# The Circadian Clock Controls Hepatic Stellate Cell Activation in Liver Fibrosis via a BMAL1/CK1ε/REV-ERBα/Transgelin Signaling Pathway

**DOI:** 10.1101/2025.09.12.675813

**Authors:** Manuel Johanns, Alexandre Berthier, Jimmy Vandel, Sandra Courquet, Julie Dubois-Chevalier, Manjula Vinod, Ninon Very, Francesco Paolo Zummo, Marie Bobowski-Gérard, Georgiana Toma, Céline Gheeraert, Didier Vertommen, Gaëtan Herinckx, Violeta Raverdy, François Pattou, Bart Staels, Jérôme Eeckhoute, Philippe Lefebvre

## Abstract

Liver fibrosis is a progressive and life-threatening condition with no effective targeted treatments. Growing evidence indicates a two-way relationship between circadian rhythm and fibrogenesis, although the specific molecular signaling pathways involved are still not well understood. The molecular clock, which governs circadian rhythms, regulates metabolic and cellular functions, and its pharmacological manipulation has shown potential as a therapy for organ fibrosis.

Although the liver’s molecular clock appeared resilient to the progression of chronic liver disease in humans from steatosis to fibrosis, detectable changes in the daily amplitude of clock genes were observed in a cohort of people living with obesity. We discovered a clock-controlled signaling pathway that drives hepatic stellate cell (HSC) activation, a key event in fibrosis progression. Interfering with this pathway, either by disrupting the core regulator CLOCK:BMAL1 or activating the nuclear receptors REV-ERBs, significantly reduced HSC activation. We also identified transgelin as the downstream effector of clock-regulated HSC contractility, a characteristic of HSC activation. Transgelin is regulated indirectly by a BMAL1-CK1ε signaling pathway and directly by REV-ERBα.

Our findings identify a previously unknown circadian-controlled mechanism that links the molecular clock to HSC activation and cell contractile function, which is relevant to human diseases. This pathway provides several entry points for drugs to target and disrupt fibrogenic signaling. By connecting clock biology to the cellular processes that cause fibrosis, our work also offers a mechanistic basis for chronotherapeutic strategies against chronic liver disease.

## INTRODUCTION

Organ fibrosis results from an excessive wound-healing response to ongoing viral, mechanical, nutritional, or other insults that remodel tissue architecture, increase stiffness, and impair organ functions. It underpins life-threatening conditions such as liver cirrhosis, pulmonary and renal fibrosis, and systemic sclerosis, for which transplantation is often the only therapy. Chronic liver diseases (CLDs) have various causes, including viral infections, excessive alcohol intake [alcohol-related liver disease (ALD)], poor diet [metabolic (dysfunction)-associated steatotic liver disease (MASLD)], or both [metabolic dysfunction and alcohol-related liver disease (MetALD)](Rinella *et al*, 2023). Liver fibrosis is a key pathological outcome of chronic liver injury and a major contributor to mortality from CLDs worldwide. It is also the most significant predictor of patient mortality from liver- or cardiovascular-related adverse events (Hagstrom *et al*, 2017; Long *et al*, 2021; Unalp-Arida & Ruhl, 2017), highlighting the importance of early detection and intervention.

Fibrosis is now recognized as a dynamic, multi-stage process triggered by tissue injury and inflammation. When this response is not resolved, it shifts from a reparative phase to a pathological one, characterized by excessive extracellular matrix (ECM) buildup. This progression occurs through a “hot fibrosis” stage, marked by active inflammation and ECM accumulation, leading to “cold fibrosis”, where ECM production dominates (de Zawadzki *et al*, 2025). In the liver, excessive ECM buildup is driven by proliferative, migratory, and contractile myofibroblasts that gather in fibrotic regions (Chung *et al*, 2022). The main source of these myofibroblasts is hepatic stellate cells (HSCs), whose transition from a quiescent to an activated state is notably accompanied by significant changes in cellular metabolism (Bates *et al*, 2020; Chen *et al*, 2012; Du *et al*, 2018; Younesi *et al*, 2024).

Cellular metabolism is controlled by exogenous cues and internal mechanisms, among which the circadian clock has emerged as a critical regulator, orchestrating the expression of hundreds of circadian output genes (Laothamatas *et al*, 2023; Schrader *et al*, 2024). The core of the circadian clock relies on a series of interlocking transcriptional-translational feedback loops (TTFLs) that generate rhythmic gene expression. At its core is a cell-autonomous TTFL where the proteins BMAL1 (encoded by the *Basic Helix-Loop-Helix ARNT Like 1* [*Arntl*] gene) and CLOCK (encoded by the *Circadian Locomotor Output Cycles Protein Kaput* [*Clock*] gene) form a heterodimer that activates the transcription of the period (*Per1*-*3*) and cryptochrome (*Cry1*-*2*) genes. PER-CRY dimers are the negative arm of the TTFL as they repress BMAL1 and/or CLOCK transcriptional activity (Smith & Sassone-Corsi, 2020). BMAL1 is considered the only essential component of this central loop (Bunger *et al*, 2000). To enhance and sustain the BMAL1 oscillation, a secondary TTFL operates in parallel in which RORα/β/γ nuclear receptors (encoded by *Rora, Rorb or Rorc* genes) act as activators and REV-ERBα and REV-ERBβ (encoded by *Nr1d1* and *Nr1d2* genes) act as repressors of *Bmal1* expression. BMAL1 itself drives the expression of both RORs and REV-ERBs, creating a “stabilizing loop” that fine-tunes the primary BMAL1:CLOCK TTFL and maintains the precision of the circadian rhythm with a period of approximately 24 hours (Cox & Takahashi, 2019). Additional layers of regulation occur through post-translational modifications of core clock proteins (Brenna & Albrecht, 2020).

Considering this, the potential link between the circadian clock, HSC activation and fibrosis has been examined through pharmacological and genetic approaches. Whole body *Bmal1* gene inactivation in *ob*/*ob* mice protects from diet-induced fibrosis (Jouffe et al, 2022), while its AAV8-mediated overexpression blunts carbon tetrachloride (CCl_4_)-induced fibrosis (Xu et al, 2022). Disruption of the BMAL1:CLOCK positive arm of the clock in ClockΔ19 mice leads to arrhythmicity and increased sensitivity to CCl_4_-induced fibrosis (Jokl et al, 2023). In contrast, *Per2*-null mice, characterized by disruption of the negative arm of the clock and subsequent increased CLOCK expression also show sensitivity to CCl_4_-induced fibrosis (Chen *et al*, 2010a; Chen *et al*, 2009). Inconsistencies among these results may stem from systemic effects and/or altered intercellular communication (Crouchet *et al*, 2025; Guan *et al*, 2020). Using REV-ERBα/β agonists to pharmacologically target the clock-stabilizing loop provides protection against fibrosis, although the proposed mechanisms vary significantly (Crouchet *et al*., 2025; Griffett *et al*, 2020; Li *et al*, 2014; Ni *et al*, 2021). Investigation of REV-ERB cell-autonomous effects by agonist treatment also supports the identification of REV-ERBs as factors promoting HSC quiescence, a conclusion further reinforced by loss-of-function studies (Chen *et al*, 2023; Crouchet *et al*., 2025; Thomes *et al*, 2016).

Reciprocally, whether pro-fibrotic conditions alter core clock gene (CCG) expression has been investigated primarily in murine model systems. A methionine and choline-deficient diet or CCl_4_ treatment, both potent inducers of liver fibrosis, disrupted the phase and amplitude of multiple CCGs (Chen *et al*., 2023; Chen *et al*, 2010b; Leclere *et al*, 2020). Although this causal relationship cannot be directly tested in humans, correlations between disruptions of rhythmic processes, such as the sleep-wake cycle, and cirrhosis have been observed (Montagnese *et al*, 2014). Mathematical extrapolation of gene expression patterns in human livers suggests that core clock components are resilient in liver cancer (Anafi *et al*, 2017), consistent with our findings in human liver biopsies with known collection times (Johanns *et al*, 2024). Additionally, biopsies from patients with different liver disease etiologies revealed a decrease in *BMAL1* expression in fibrosis, along with decreased expression of circadian clock repressors (Jouffe *et al*., 2022).

In this study, we conducted a comprehensive examination of CCGs expression in livers of a time-of-day-controlled human metabolic dysfunction-associated steatohepatitis (MASH) and fibrosis cohort. While time-of-day dependent CCG expression remained largely resilient, we identified significant alterations in its amplitude among patients with MASH and fibrosis. In mice, profiling of CCG expression in liver parenchymal and non-parenchymal cells demonstrated the exquisite sensitivity of the molecular clock in hepatic stellate cells (HSCs) and cholangiocytes (CHs) to both acute and prolonged CCl₄ administration. Mechanistically, the CLOCK:BMAL1/REV-ERB axis was implicated in mediating the pro-fibrotic response in primary human HSCs and the immortalized mouse HSC line, EMS404 which exhibited robust circadian rhythmicity and responded to transforming growth factor beta (TGFβ) stimulation. In this context, REV-ERB agonism and BMAL1 inhibition effectively abrogated TGFβ-induced HSC activation and reduced cell contractility without disrupting the canonical TGFβ signaling pathway. These effects were corroborated at the transcriptional level by a pronounced suppression of the pro-fibrotic program, as evidenced by the downregulation of ECM- and focal adhesion-related gene expression. Finally, a combined transcriptomic and proteomic approach identified transgelin as a molecular relay for CCG-mediated regulation of fibrosis, suggesting transgelin inhibition as a novel therapeutic strategy against hepatic fibrosis.

## MATERIALS AND METHODS

### Reagents

SR9011 (#HY-16988), KL001 (#HY-108468), CLK8 (#HY-148765) and SB431542 (#HY-10431) were purchased from MedChemExpres. Recombinant human TGFβ1 (#240-B-002) was from R&D Systems.

Anti-SMAD2/3 (#5678), anti-phospho-SMAD2 (Ser465/467)/SMAD3 (Ser423/425) (#8828), anti-BMAL1 (#14020), anti-TAGLN (#36090), anti-PhosphoGSK3 #5558) and anti-CLOCK (#5157) antibodies were from Cell Signaling Technology. Anti-COL1A1 (#ab34710) and anti-αSMA (ACTA2) (#ab124964) antibodies were from Abcam. The anti-REV-ERBα antibody (#A8740A) was from Perseus. The anti-HSP90AA1 antibody (#662802) was from Biolegend. HRP-conjugated anti-rabbit (#A4416) or anti-mouse (#A0545) IgGs were from Sigma.

Fluorochrome-coupled antibodies targeting CD31-BV421 (#BLE102424), CD45-BV510 (#BLE103138), CD326-CF594 (#BLE118236) F4/80-PE-Cy7 (#BLE123114) CD146-APC (#BLE134712) MHCII-AF700-(#BLE107622) CD11b-APC-Cy7 (#BLE101226) were purchased from BioLegend. Anti-CLEC4F was from R&D (#MAB2784) and coupled to CF568 using the Mix-n-Stain CF568 Antibody Labeling Kit (Biotium, #BTM92235).

### The HUL/ABOS cohort

The Hôpital Universitaire de Lille (HUL) cohort, also referred to as the Biological Atlas of Severe Obesity (ABOS; ClinicalTrials.gov: NCT01129297), was established in 2006 at the University Hospital of Lille, France, to recruit severely and morbidly obese patients eligible for bariatric surgery. The study protocol adheres to the ethical principles of the 1975 Declaration of Helsinki, and written informed consent was obtained from all participants. Patients had to fast from midnight until surgery. During the surgical procedure, liver wedge biopsies were collected, immediately snap-frozen, and the exact time-of-day of collection was recorded. To date, the cohort comprises over 2,000 patients, from whom >900 liver biopsies were histologically and transcriptionally characterized (Vandel *et al*, 2021). From a 319-patient cohort with clear histological features, a propensity-matched (on age, BMI, insulin resistance, and fibrosis) sub-cohort was defined, while keeping a constant male:female ratio, an equal number for each disease state, and equal distributions between samples collected in the morning (AM) or afternoon (PM). This yielded a 213-patient cohort comprised of 71 (histologically normal, NL), 71 steatosis and 71 MASH cases. Main clinical, histological and molecular characteristics are summarized in **Supp.** Figure 1A, with statistical analyses performed using the “gtsummary” package (v1.5.0). The “F0 vs >F2” subcohort has been described elsewhere (Bobowski-Gerard *et al*, 2022).

### Animal experimentation

Animals were handled following institutional guidelines and approved by Comité d’Ethique en Expérimentation Animale n°075, Lille, France. Eight to 12-week-old C57Bl/6J male mice were purchased from Charles River Laboratories. Per2::Luc mice were from The Jackson Laboratory [IMSR_JAX:006852, (Yoo *et al*, 2004)] and were rederivated into Specific Opportunist Pathogen Free C57Bl/6J mice at Charles River Laboratories. Mice were housed in a 12h/12h light/dark cycle at 22-24°C for 2 weeks before experimentation. Animals were fed ad libitum on a chow diet (Safe diets, A04) with free access to drinking water. Liver fibrosis was initiated by intraperitoneal injection as indicated (single injection or 3 times a week for 2 weeks with 0.5 mL/kg CCl_4_ (#319961, Sigma-Aldrich) diluted in olive oil). The last injection of CCl_4_ was done 24 h before organ collection.

### Liver parameters

Blood alanine aminotransferase (ALT) and aspartate aminotransferase (AST) activities were measured with colorimetric assays (#981769 and 981771, Thermo Fisher Scientific Inc., MA, USA) using a Konelab 20 Clinical Chemistry Analyzer (Thermo Fisher Scientific Inc., MA, USA).

### Liver explant preparation and ex vivo luciferase activity monitoring

A detailed protocol is described elsewhere (Berthier et al, 2025). Briefly, Per2::luc mice were sacrificed by cervical dislocation at ZT7 (7 h after lights-on). Livers were immediately placed in ice-cold Hanks’ balanced salt solution, and the left lobe was sectioned into 2 mm³ pieces. Fifteen explants were positioned on a Millicell insert (Millipore #PICMORG50) in a 35 mm dish containing bicarbonate-free Dulbecco’s modified Eagle medium (Sigma #D5030) supplemented with 10 mM HEPES (pH 7.4), 25 mM glucose, 10% fetal calf serum, 4 mM sodium bicarbonate, 1% Glutamax, 1% penicillin-streptomycin, and 200 µM beetle luciferin. Explants were cultured at 36°C without CO₂ and bioluminescence was recorded (1 min every 10 min) using a KRONOS-DIO AB-2550 system. Data were analyzed with ATTO Kronos software (v2.30.243).

### Liver cell type isolation

A detailed protocol is described elsewhere (Zummo et al, 2023). Briefly, parenchymal and non-parenchymal liver cells were obtained from male C57Bl6/J mice (12–17 weeks, Charles River). After euthanasia by cervical dislocation, livers were perfused and dissociated with collagenase IV (100 U/mL, Sigma). Hepatocytes were collected by centrifugation and the supernatant incubated with collagenase, pronase (0.5 mg/mL), and DNase I (10 µg/mL) at 37°C. All subsequent steps were performed at 4°C in the dark. After red blood cell lysis, 2–3×10⁶ cells per tube were stained with Zombie Green (Biolegend) and incubated with BD FcBlock (anti-CD16/32, BD #553142), followed by antibody staining (20 min). Antibodies targeting CD31, CD45, CD326, F4/80, CD146, MHCII, CD11b (BioLegend) and CLEC4F (R&D Systems, labeled with CF568, Biotium) were used to distinguish between hepatic stellate cells, liver sinusoidal endothelial cells, cholangiocytes and Kupffer cells. Sorting was performed on a BD INFLUX v7 cell sorter (BD Biosciences). Data were analyzed with FlowJo (v10.5.3). Sorted cells were collected in RNAlater (ThermoFisher) for RNA extraction.

### Sirius red staining

The liver median lobe was fixed in 4% paraformaldehyde for 48 h, paraffin-embedded, and sectioned at 5 μm onto gelatin-coated slides. Fibrosis was assessed as described (Very *et al*, 2024). Briefly, liver sections were stained with 0.1% Sirius Red in 1.3% picric acid. Entire sections were scanned (Axioscan Z1 slide scanner, Zeiss), and fibrosis quantification was performed on 10 randomly selected fields using ImageJ excluding vessel-containing areas.

### RNA extraction and RT-QPCR

RNAs from liver was extracted as follows. Fifty mg liver samples were homogenized using a Polytron in Trizol (Invitrogen, #15596026) and after addition of chloroform (1:5, v:v), samples were centrifuged for 5 min at 4°C, 15,000 x g. Upper phases were collected and supplemented with 3 volumes 100% ethanol. RNAs were purified using Nucleospin RNA columns (Macherey-Nagel, #740955.250S) following the manufacturer’s recommendations.

Cellular RNAs were extracted from cells using the Macherey-Nagel™ Nucleospin™ Mini kit (#872061) or Qiagen RNeasy Micro kit (#74004), depending on sample abundance, following the manufacturers’ protocols. RNA concentration and purity were assessed with a Nanodrop One (ThermoFisher Scientific) or Qubit fluorometer and RNA HS Assay kit (#Q32852). RNA integrity was evaluated using a Bioanalyzer 2100 (Agilent). Samples with RIN <6.0 were discarded. RNA was reverse-transcribed with random primers using the High Capacity cDNA Reverse Transcription Kit (ThermoFisher/Applied Biosystems, #4368814). Quantitative PCR was performed in technical triplicates from at least three independent biological replicates using the SYBR Green Brilliant II Fast kit (Agilent Technologies) on a QuantStudio 3 (Applied Biosystems). mRNA levels were normalized to Rps28 and Rplp0 and fold changes calculated using the 2^−ΔΔCt^ method (Schmittgen & Livak, 2008). Primer sequences are listed in **Supplemental Table 1**. Primer efficiency was verified by serial cDNA dilution (1- to 500-fold), amplicon size by agarose gel electrophoresis, and specificity by melting curve analysis.

### Primary human HSCs

Cryopreserved primary hepatic stellate cells from a healthy donor (donor A, male) were from Innoprot (P10653). Donor B HSCs were isolated from a cirrhotic male donor at HUL (Bobowski-Gerard *et al*., 2022). Both primary HSCs were grown in DMEM supplemented with 15% FCS. Cells were treated at 70-80% confluency (passage 8-10) with DMSO (control) or 5 µM SR9011 or 5 µM CLK8 for 24 h, then with 5 ng/mL TGFβ or vehicle in the same media for an additional 24 h.

### Cell lines, plasmids and lentiviruses

Murine EMS404 cells were from Kerafast (#EMS404) and initially described in (Guo et al, 2009). EMS404 cells were cultured in Dulbecco’s Modified Eagle Medium (DMEM) supplemented with 10% fetal bovine serum (Dutscher) and 100 U/mL penicillin and 100 µg/mL streptomycin.

EMS404-BmalLuc cells were generated as follows: pLV6-Bmal-luc was a gift from Steven Brown and obtained from the Addgene collection (plasmid #68833). pLV6-Bmal-luc was transfected using JetPEI (#101000053, Polyplus) in 293T-N HEK cells together with pRSV-Rev, pMDLg/pRRE, pMD2.G plasmids encoding HIV Rev, viral antigens (ag), the reverse transcriptase *Pol* (GAG-Pol), and the VSV-G envelope glycoprotein (VSV-G), respectively. Culture supernatants containing lentiviral particles were collected 48 h later, filtered on a 0.45 µm filter and concentrated using an AmiconTM ultrafiltration unit with a 100 kDa-cutoff before spreading on EMS404 cells at 70% confluency. Blasticidin (#SBR00022, Sigma-Aldrich)-resistant cells were selected and verified for bioluminescence in the presence of luciferin.

REV-ERBα-depleted EMS404 cells were generated as above using pLKO-1 shRev-Erbalpha, a gift from Bruce Spiegelman [Addgene plasmid #22747;(Estall et al, 2009)].

EMS404 cells overexpressing REV-ERBα (EMS LV-FlagNr1d1) were generated by transduction with lentiviral particles containing a FLAG epitope-tagged version of human *NR1D1* (Berthier et al, 2018). The flagged h*NR1D1* ORF was inserted into the pLenti-CMV-GFP-Puro (658-5) backbone engineered to remove the GFP ORF. pLenti-CMV-GFP-Puro (658-5) was a gift from Eric Campeau & Paul Kaufman [Addgene plasmid #17448;(Campeau et al, 2009)].

Control EMS404 cells (LV-empty) were generated using the pLKO.1 puro vector, a gift from Bob Weinberg [Addgene plasmid #8453; (Stewart et al, 2003)].

### Small interfering RNA transfection

Scrambled control siRNAs (Smartpool #D-001810-10-05), siRNAs targeting mouse *Bmal1* or *Tagln* transcripts (L-040483-01-0005 and L-062786-01-005, respectively) were from Dharmacon. siRNAs were used at 50nM final concentration and transfected with Dharmafect1 transfection reagent (#-2001-01) in antibiotic-free DMEM. EMS404 cells were seeded in 24-well plates and transfected at around 70% confluency according to the manufacturer’s instructions. Six hours after transfection, the medium was replaced with fresh, serum-supplemented DMEM. Twenty-four hours after transfection, TGFβ1 was added as indicated. Cells were harvested 48 h after transfection.

### Collagen gel contraction assay

Collagen (from rat tail tendon, #11179179001, Roche Diagnostics GmbH) was dissolved in filter-sterilized 0.2% acetic acid at a concentration of 3 mg/mL. An EMS404 cell suspension (5 × 10⁶ cells/mL, passages 2–7) was prepared in 1× DMEM (#D2429, Sigma) and mixed immediately with the NaOH-neutralized collagen solution to achieve a final cell density of 1.25 × 10⁶ cells/mL. The mixture was then distributed into 6-well plates. Following polymerization of the collagen lattice, the medium was added on top and the indicated treatments were applied. On day 1, the collagen disc was gently detached from the bottom of the well. The surface area of the collagen disc was measured daily until day 5 and analyzed with ImageJ.

### Seahorse assays

The glycolytic rate of EMS404 cells was measured in real time using an XF96 Extracellular Flux Analyzer (Agilent Technologies, Santa Clara, CA, USA). Cells were seeded at a density of 75,000 cells per well in an XFe96/XF Pro Cell Culture Microplate (#1037-94-100, Agilent) with standard culture medium. After 24 h, cells were washed with 1× PBS (Corning) and treated with TGFβ (1 ng/mL or 5 ng/mL) for an additional 2 h. Following treatment, the culture medium was replaced with assay medium composed of XF DMEM (pH 7.4; #103575-100, Agilent) supplemented with 1 mM pyruvate (#11360-039, Gibco), 2 mM glutamate (#35050-038, Gibco), and 25 mM glucose (#25-037-CI, Corning). Cells were incubated in this medium for 45 min to allow acclimatization before measurement. Injection solutions were prepared in the same assay medium according to the manufacturer’s instructions for the Glycolytic Rate Assay. Specifically, rotenone (50 mM; #282T2970, TargetMol Chemicals, Boston, MA, USA), antimycin A (5 mM; #J63522, ThermoFisher), and 2-deoxy-glucose (500 mM; #D6134-5G, Sigma-Aldrich) were loaded into the designated ports of the XFe96/XF Pro sensor cartridge (#103793-100, Agilent). The sensor cartridge and cell plate were then placed in the flux analyzer, and the Glycolytic Rate Assay protocol was initiated. Data were collected as the extracellular acidification rate (ECAR), which served as an indicator of glycolytic activity. Differences in glycolytic rate between conditions were analyzed based on ECAR measurements.

### RNA sequencing, data processing and analysis

After initial QCs (OD260/280, RIN), oligo dT selection, random (N6) primed reverse transcription, end repair and adaptator ligation, cDNAs samples were paired-end sequenced (100 b) on the DNBSEQ BGI platform. Sequence data filtering was performed using BGI SOAPnuke (v 1.5.2)(parameters: -I 15 -q 0.2 -n 0.05) and cleaned data were stored as a Fastq file. HISAT [v 2.0.4, (Kim et al, 2015)] was used to align valid reads on the mouse genome (mm10)(parameters:--sensitive –no-discordant –no mixed -I 1 -X 1000 -p8 –rnastrandness RF). Bowtie2 [v2.2.5, (Langmead & Salzberg, 2012)](parameters: -q –phred64 –sensitive –dpad0 –gbar99999999 –mp1,1 -np 1 –score-min L,0,-0.1 -p 16 -k 200) was used to map reads of the reference transcriptome. RSEM was used to determine gene expression levels [v1.2.8, (Li & Dewey, 2011)]. DEseq2 was used to identify differentially expressed genes [q value < 0.05, (Love et al, 2014)].

### ChIP-Seq

ChIP-seq data were reprocessed as described in (Dubois-Chevalier et al, 2017) from GSE39977 [BMAL1 cistrome, (Koike et al, 2012)] and GSE26345 [REV-ERBα cistrome, (Bugge et al, 2012)]. Tracks were visualized using IGV [v2.17.2.02 (Robinson et al, 2011)].

### WES analysis

Protein Simple Wes analysis was performed as previously described (Vinod *et al*, 2022).

### Proteomic analysis-Tandem Mass Tag isobaric labeling

EMS404 cells were seeded in 10 cm culture dishes at approximately 60% confluency. After 24 h, the medium was replaced with DMEM supplemented with 0.5% FBS and 0.2% BSA.

On day 4, cells were washed twice with ice-cold 1× PBS and lysed directly in the dish with 300 µL of lysis buffer (50 mM triethylammonium bicarbonate, pH 8.5 [#90114, Thermo Fisher], 150 mM NaCl, 1% Igepal, 0.5% sodium deoxycholate, 0.2% n-dodecyl-β-D-maltoside, and 0.1% SDS), freshly supplemented with 10 mM NaF, 1 mM sodium orthovanadate, 1 mM PMSF, 2.5 mM β-glycerophosphate, and 1 mM DTT. Lysates were harvested by scraping, vortexed, incubated on ice for 10 min, vortexed again, and centrifuged at 14,000 × g for 10 min at 4 °C. Protein concentrations were determined using the BCA assay.

A total of 500 µg of protein was transferred to low-protein-binding tubes. Proteins were precipitated by adding four volumes (1 mL) of pre-chilled acetone (−20 °C) and incubating overnight at −20 °C. The resulting pellets were recovered by centrifugation (2,000 × g, 5 min, 4 °C), washed 3 times with 1 mL of cold acetone (with vigorous vortexing), air-dried under a chemical hood for 10–15 min, and resuspended in 100 µL of 100 mM TEAB (pH 8.5).

Samples were digested overnight at 37 °C using sequencing-grade trypsin (#V5111, Promega) at an enzyme-to-substrate ratio of 1:100. Specifically, 20 µL of 0.2 µg/µL trypsin was added per 100 µL sample (5 µg trypsin for 500 µg protein).

Following digestion, samples were thoroughly dried using a SpeedVac concentrator (≥2 h), and either stored frozen or processed for downstream mass spectrometry analysis. Peptides were quantified using the Thermo Pierce Peptide Assay (#23275, Thermo Scientific), and 10 µg of each sample was labeled with the TMTpro 16plex Labeling Kit (#A44521, Thermo Scientific). Labeled samples were pooled and fractionated using the high-pH reverse-phase peptide fractionation kit (#84868, Thermo Scientific). Peptide fractions were dried, resuspended, and 0.8 µg of each was analyzed using a Fusion Lumos Orbitrap mass spectrometer.

Protein abundance data were normalized and scaled using the Proteome Discoverer software (v.2.5, ThermoScientific). Peptides were filtered out if Sum.PEP.Score <5. More than 5,000 known proteins were identified according to this analysis pipeline (Jacobs *et al*, 2025).

### Human transcriptomic data analysis

Transcriptomic data collection and initial processing were described in (Vandel *et al*., 2021). To achieve statistical power and obtain robust and exhaustive lists of time-dependent gene expression over the available daytime window, 3 complementary statistical methods analyzing different aspects of gene expression distribution (differential expression, partial Spearman correlation, Kolmogorov-Smirnov test) were used. The results of these analysis were agglomerated by a Fisher test to yield a combined p value for each gene. A detailed protocol is available at (Johanns et al., 2024). Briefly, analyses were performed in R (v4.1.0) using RStudio (v1.4.1106). Clinical data (**Figure 1D**) were summarized using the gtsummary package (v1.5.0). For transcriptomic analyses, RSEM transcript counts were rounded and summed per gene, then pre-filtered to exclude genes with total counts below the number of samples (i.e. mean <1 read per sample). Differential expression between morning (AM, before 12:00) and afternoon (PM, after 12:00) biopsies was assessed using DESeq2 (v1.43.0), with sex included as a covariate in the model.

**Figure 1.**
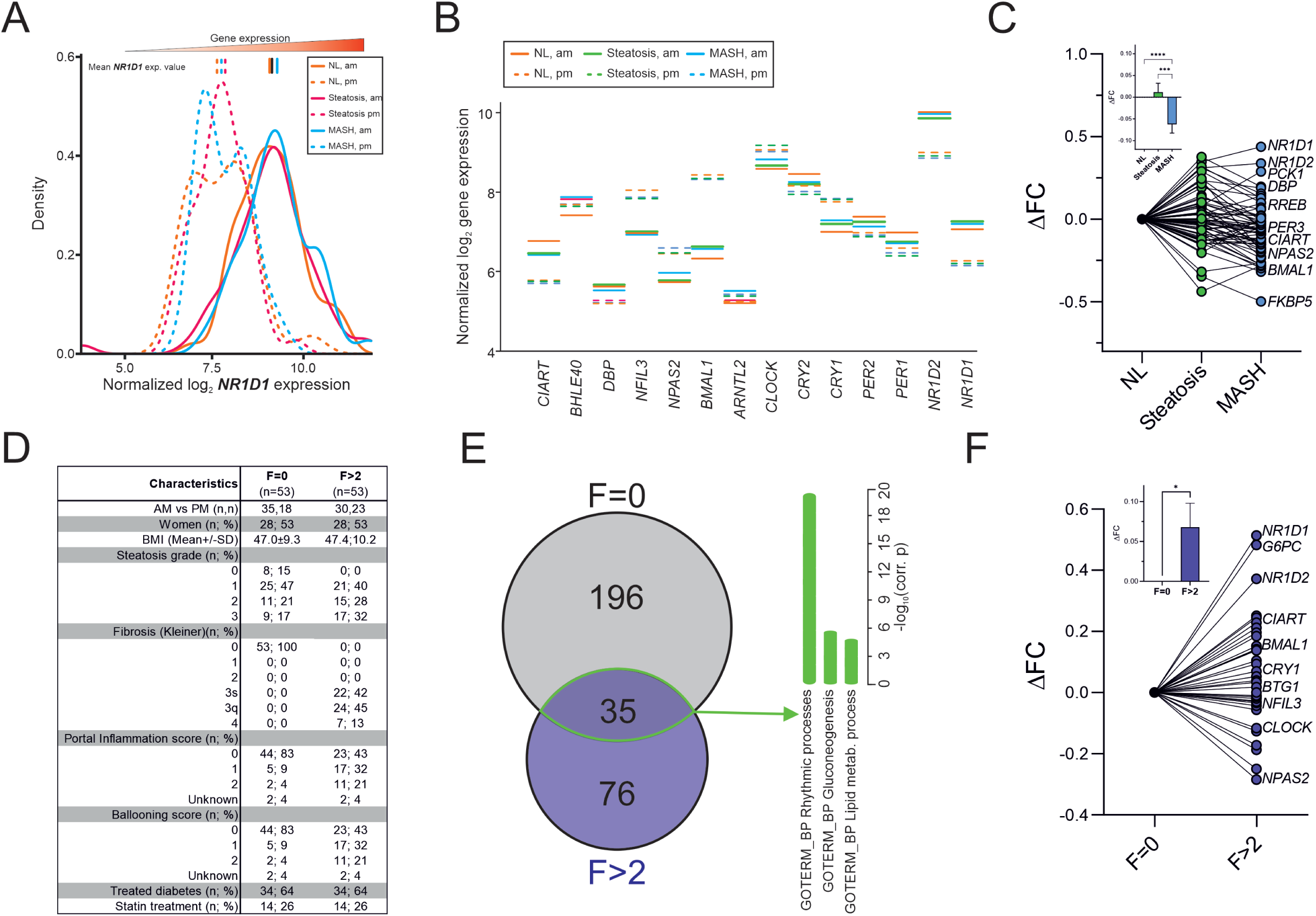
Transcriptomic analysis of core clock genes in MAFLD and fibrosis. A) CCG frequency distribution in human livers. Relative frequency distribution of *NR1D1* expression values in normal (NL), steatotic and MASH liver biopsies taken in the morning (AM; continuous lines) *vs.* afternoon (PM; dotted lines). Group mean values are indicated as corresponding vertical lines at the top of the graph. B) CCG mean expression in human liver. Mean expression values for core clock genes (CCGs) in normal (NL), steatotic and MASH liver biopsies taken in the morning (AM; continuous lines) *vs.* afternoon (PM; dotted lines). C) CCG amplitude in human livers. Difference in fold changes (ΔFC) between mean AM and PM expression of common daytime-dependent genes in steatotic and MASH livers, compared to those of normal liver (NL). *Inset:* Group means ± s.e.m. of fold changes. D) Characteristics of the subcohort. Biometric, biochemical, and histological parameters of propensity score-matched control (Kleiner fibrosis score F = 0; n=53) and fibrotic (F > 2; n=53) patients. E) TDG gene distribution in fibrosis. Overlap of daytime-dependent genes identified by RNAseq in fibrotic (F>2) *vs.* non-fibrotic (F=0) liver, along with enriched functional GO-BP terms. F) TDG amplitudes in human fibrosis. ΔFC between mean AM and PM expression of common daytime-dependent genes in fibrotic (F>2) compared to non-fibrotic (F=0) livers. Values were compared using an unpaired two-tailed t-test, with p<0.05 judged significant.

To identify robust time-dependent genes, 2 additional approaches were applied to variance-stabilized counts: (1) a Kolmogorov-Smirnov test (stats, v4.1.0) comparing AM vs. PM distributions, and (2) a partial Spearman correlation with time-of-day adjusted for sex using ppcor (v1.1). P-values from the three methods were combined per gene within each NAFLD group using Fisher’s method (metap, v1.7) and adjusted for multiple testing using Benjamini-Hochberg correction (stats, v4.1.0). Genes were defined as time-dependent if the adjusted Fisher p-value (FDR) was <0.01, optionally with an absolute fold-change >1.2 (Johanns et al., 2024).

### Statistical analysis

Except for omics data, statistical analyses were conducted using GraphPad Prism (v10.4.1 or earlier). Data are presented as mean ± standard error of the mean (s.e.m.), with a minimum of 2 independent biological replicates per experiment. For in vitro datasets, variance equality was assessed using the F-test. Comparisons between 2 groups were performed using an unpaired two-tailed t-test with Welch’s correction. For multiple group comparisons involving a single variable, a one-way ANOVA followed by Tukey’s post hoc test (comparing all groups pairwise) was applied. When more than 1 variable was involved, a two-way ANOVA followed by Tukey’s multiple comparisons test was used. A p-value of less than 0.05 was considered statistically significant. All data represent biological, not technical, replicates.

### Data visualization

Generally, graphs were generated with GraphPad Prism (version 10.4.1 or earlier). Bubbleplots were either generated in R studio using the ggplot2, plotly, reshape2, rcpp and tidyverse packages, or through the proprietary BGI’s Dr Tom interface. SVG files were modified with Inkscape v1.3.2 and assembled as figures using CorelDRAW 2020.

### Data availability

RNA-seq datasets are available at NCBI GEO under the accession numbers GSE304664 and GSE304977.

## RESULTS

### Time-of-day differential expression of core clock genes is altered in steatotic and MASH livers

From an initial cohort of 620 patients living with obesity and with complete clinical, biometric, histological data as well as recorded time-of-day of biopsy collection and liver transcriptomic data (Vandel *et al*., 2021), we constructed 3 equally sized groups (n = 71 each; 213 patients in total) with a consistent male-to-female ratio. These groups were optimally matched for age, BMI, insulin resistance (HOMA-IR), and fibrosis stage to minimize potential bias and control for known confounders. Each group represented one liver condition: healthy (NL, normal liver), steatotic, or MASH (**Supp.** Figure 1A). A multivariate analysis of variance indicated, as previously reported (Johanns *et al*., 2024), that sex, histology, and time of day were significant explanatory variables, as supported by the corresponding F ratios (**Supp.** Figure 1B). Biopsy collection times were similarly distributed across groups, with most samples (60-65%) collected in the morning (AM) with a peak collection time around 9:30 AM. Afternoon (PM) samples, with a collection peak at 1:30 PM, accounted for approximately 35–40% of the total. (**Supp.** Figure 1C). We used our combinatorial statistical method combining 3 different analysis of gene expression distribution (differential expression, partial Spearman correlation, and Kolmogorov-Smirnov test) aggregated by a Fisher test (Johanns *et al*., 2024) to identify significantly time-dependent genes (TDG) for each group. As previously reported, many TDGs were specific to each group (44%-64%), with a core conserved set of genes (12%, 66 out of 552 total) whose functions are associated with circadian rhythmicity and glucose and lipid homeostasis (**Supp.** Figure 1D). Among these genes were CCGs (**Supp. Table 2**), thus confirming our initial analysis carried out on a broader cohort (Johanns *et al*., 2024). We calculated the mean expression value for each CCG and compared AM vs. PM values across groups (**Figure 1A, B**). Fold change variation analysis across groups revealed that AM to PM amplitudes (ΔFC) mildly increased in steatosis, and globally decreased in MASH livers (**Figure 1C, inset**), with *NR1D1* and *NR2D2* showing increased amplitudes while BMAL1 amplitude was reduced (**Figure 1C**).

A similar approach was followed to compare non-fibrotic (Kleiner score F=0, n=53) to matched fibrotic (Kleiner score F>2, n=53) livers (**Figure 1D**). Common TDGs included again CCGs (*NR1D1*, *NR1D2*, *NPAS2*, *CIART*, *PER3*, *CLOCK*, *BMAL1*, *CRY1*, *NFIL3*)(**Figure 1E**), yet for some of them, the AM vs. PM differential expression was altered in fibrosis (**Figure 1F**). Fold change variation analysis across groups revealed that AM to PM amplitudes globally increased in fibrotic livers (**Figure 1F, inset**). Individually, *NR1D1*, *NR2D2* and *BMAL1* transcripts showed increased AM vs PM amplitudes, whereas the BMAL1 dimerization partner *CLOCK* displayed a lower amplitude in fibrosis.

Taken together, our data from human livers indicate that although CCGs remain resilient in the context of MASH and fibrosis, they exhibit detectable alterations in their time-of-day-dependent amplitude across conditions. However, in the absence of complete circadian expression profiles, and given interindividual variability and the variability in biopsy collection times, we cannot determine whether these changes reflect alterations in overall expression levels, amplitude, period, or a combination thereof. These observations prompted us to further investigate the behavior and role of CCGs in mouse models of fibrosis.

### Fibrosis progression in mouse liver perturbs core clock gene expression in several cell types

The hepatotoxicant CCl_4_ was used in C57Bl6/J male mice to induce a pericentral injury and a profibrotic response in the liver (**Figure 2**). Twenty-four hours after the first injection, hepatocytes (HC), stellate cells (HSCs), Kupffer cells (KCs), sinusoidal endothelial cells (LSECs) and cholangiocytes (CHs) were isolated (around ZT3, i.e. 3 hours after lights-on) and mRNA levels of selected fibrosis markers (*Acta2*, *Col3a1*, *Col1a1*, *Dpt…*) were evaluated (**Figure 2A**). *Acta2* and *Ankrd1* were strongly induced in HSCs, while *Tgfb1* expression increased in KCs. After a 2-week CCl_4_ treatment, *Acta2*, *Col1a1* and *Dpt* expression were significantly increased in HSCs while *Tgfb1* expression returned to normal in KCs (**Figure 2B**). Liver histology and serum markers of liver injury clearly indicated established fibrosis under this condition (**Figure 2C**). Since this suggested that the profibrotic milieu might evolve, we monitored the expression level of the most representative CCGs in each condition (**Figure 2D**). Interestingly, CCG expression was differentially affected by both treatment duration and cell type. The most pronounced changes were observed in HSCs, while in KCs CCGs remained largely unaffected. Most CCGs showed a decreased expression in response to CCl_4_, a trend that was also confirmed by transcript analysis from bulk liver tissue following long-term CCl₄ exposure (8 weeks, **Figure 2E**).

**Figure 2:**
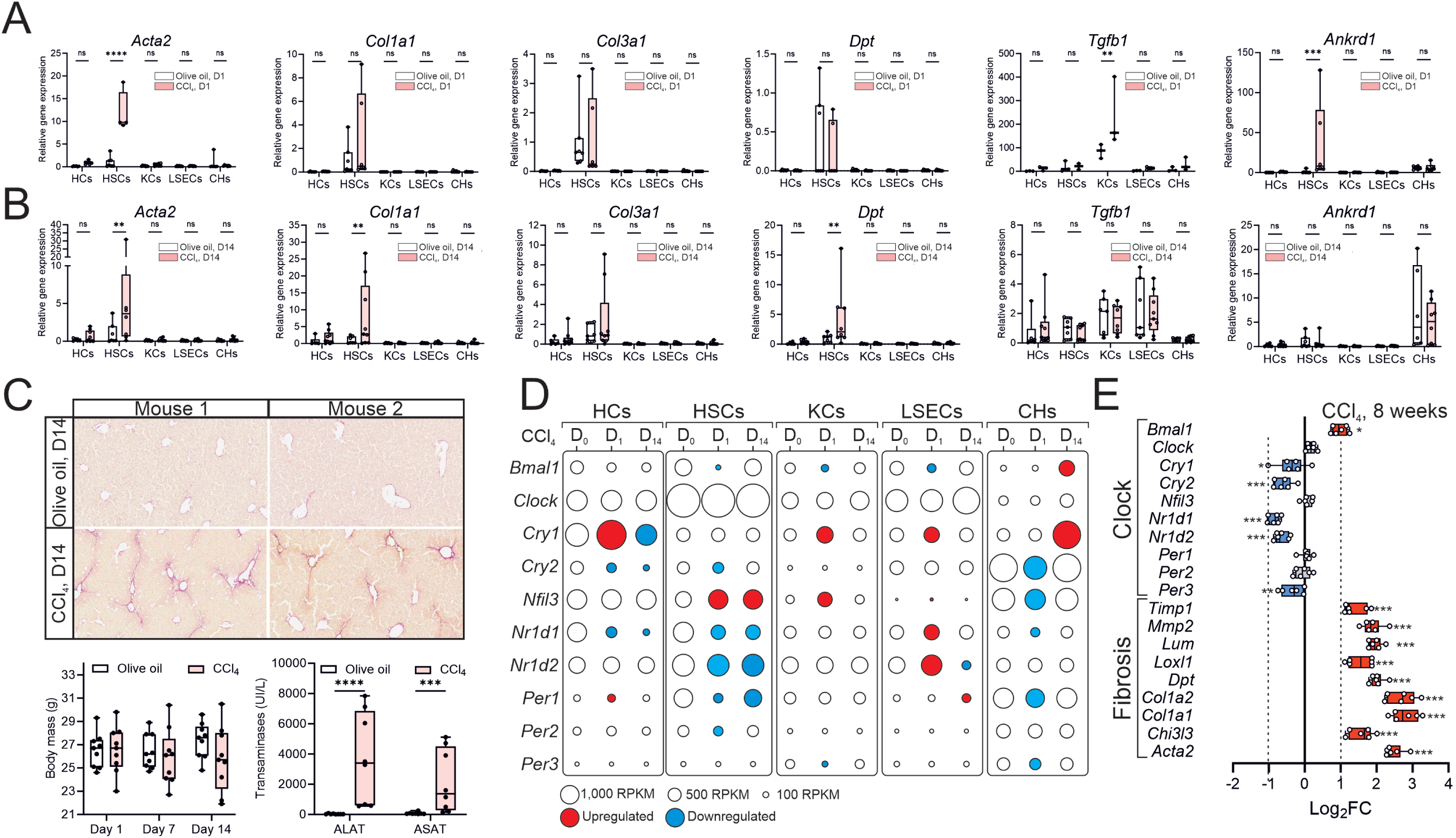
Core clock gene expression in a mouse model for liver fibrosis. A, B) Kinetics of profibrotic gene expression in different liver cell types. Hepatocytes (HCs), hepatic stellate cells (HSCs), Kupffer cells (KCs), sinusoidal cells (LSECs) and cholangiocytes (CHs) were isolated from mice injected with a single dose (24 h, A) or six doses spaced by 2-3 days (2 weeks, B) of CCl_4_ (0.5 mL/kg bodyweight). Gene expression was assayed by RT-qPCR. Averaged expression values (mean ± S.E.M., n=4-6 per cell type) were compared using an unpaired two-tailed t-test. *: p<0.05, **: p <0.01, ***: p<0.005, ****: p<0.001. C) Fibrosis assessment. Livers were collected after a 2 week-treatment with CCl_4_ and liver sections were stained with Sirius Red (upper panel) to assess collagen fiber deposition. Body mass was monitored weekly and serum alanine/aspartate transaminase (ALAT/ASAT) activities were measured as indicators of liver injury (lower panel). Averaged values (mean ± S.E.M., n=7-9 animals/group) were compared an unpaired two-tailed t-test. ***: p<0.005, ****: p<0.001. D) CCG expression in liver cell types. Core clock gene (CCG) expression by RNAseq in sorted liver cell types. Red circles indicate upregulated genes, blue circles indicate downregulated genes (one-way ANOVA, q<0.05). E) CCG expression after prolonged exposure to CCl_4_. CCG gene expression was measured by RT-qPCR in bulk liver harvested at ZT6-8, after an 8-week CCl_4_ treatment (0.5 mL/kg, 3x per week). Values are relative expression values compared to vehicle (olive oil) treatment from 6 mice per group. Statistical analysis was done using an unpaired two-tailed t-test (*: p<0.05, **: p <0.01, ***: p<0.005, ****: p<0.001).

Taken together, these data suggest that despite an evolving fibrotic milieu, CCG expression is mostly downregulated in HSCs and to a lesser extent in CHs.

### Chemical interference with the molecular clock attenuates the ex vivo pro-fibrotic response

Since *Cry2*, *Nr1d1*, *Nr1d2* and *Bmal1* expression was altered in CCl_4_-activated HSCs, the ability of pharmacological modulators targeting CRYs, REV-ERBs or BMAL1 to interfere with the liver fibrotic response was interrogated in an ex vivo model of fibrosis (**Figure 3A**). Liver explants from Per2::Luc mice (Yoo *et al*., 2004) exhibit a bioluminescent rhythm with a ≈24-hour period (**Figure 3B**) and develop a spontaneous fibrotic response that peaks after 6 days in culture (**Figure 3C**, compare D_0_ vs DMSO). The REV-ERBα/β agonist SR9011 (Solt *et al*, 2012), the BMAL1:CLOCK disruptor CLK8 (Doruk *et al*, 2020) and the CRY stabilizer KL001 (Hirota *et al*, 2012) effectively perturbed the circadian activity of the Per2-Luc fusion protein, substantiating their efficacy in this model system (**Figure 3B**). When monitoring the expression of archetypical fibrotic players (*Acta2*, *Col1a1*, *Col3a1*, *Timp1*) in the presence of increasing doses of these compounds, both CLK8 and SR9011 efficiently blunted the induction of these genes in cultured liver explants at Day 6, whereas KL001 was less effective (**Figure 3C**) and was therefore not investigated further.

**Figure 3:**
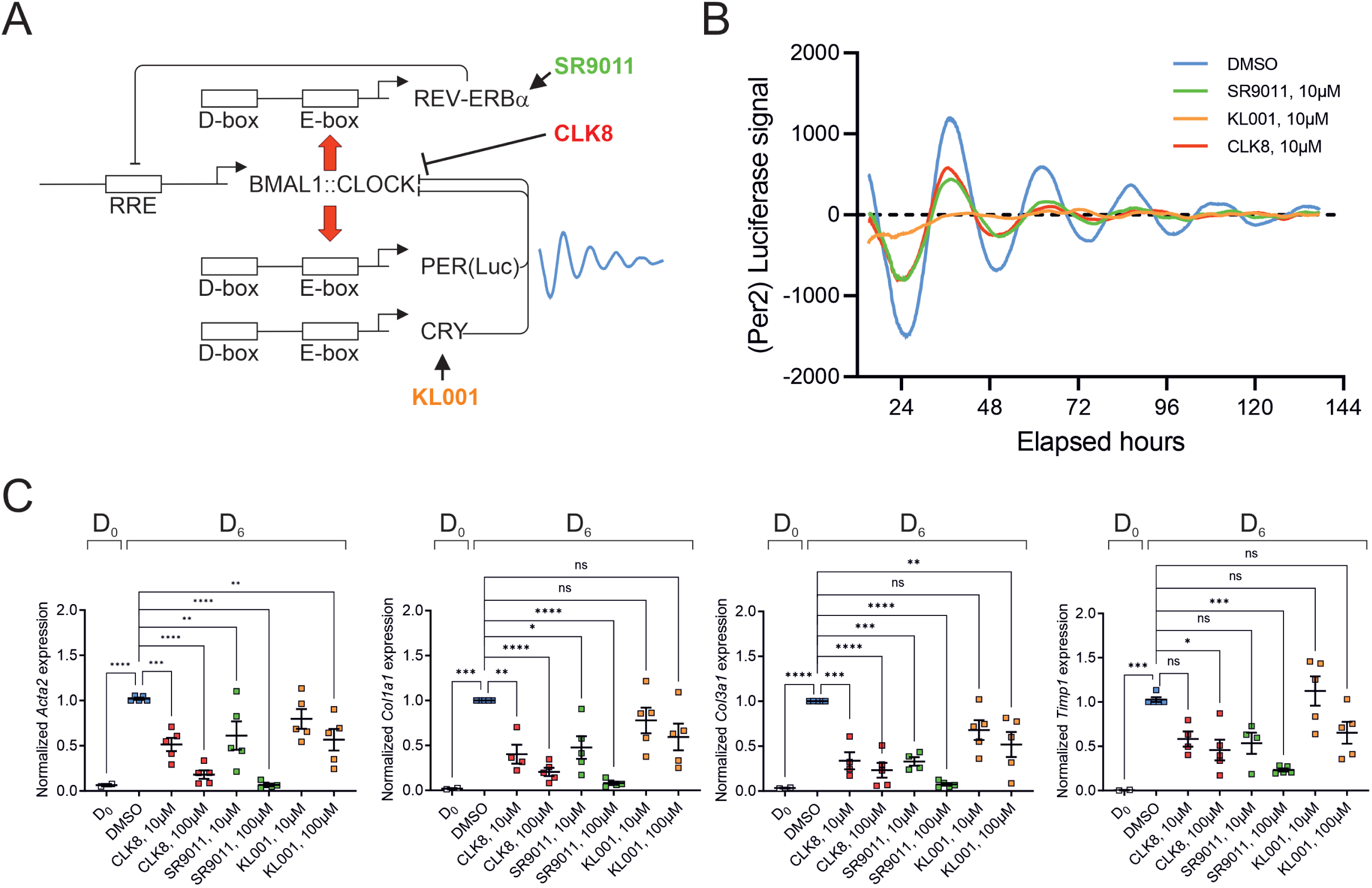
Pharmacological targeting of core clock proteins in a mouse explant model of spontaneous fibrosis. A) Schematic representation of the main core clock TTFLs and of the experimental system. B) Circadian rhythmicity in mouse liver explants. Liver explants were prepared from Per2::Luc mice bearing a luciferase fused to the endogenous core clock gene Per2. Explants were cultured in the presence of the indicated treatments and after an adaptation period of 24 h, bioluminescence was recorded for 6 days. Data are detrended from one representative experiment. C) Profibrotic gene expression in cultured liver explants. Profibrotic gene expression levels were measured by RT-qPCR in Per2::Luc mouse liver explants immediately after isolation (D0) or after 6 days (D6) in culture in the presence of indicated treatments. Averaged expression values (mean ± S.E.M., n=5) were compared for each gene using one-way ANOVA followed by Tukey’s post hoc test. *: p<0.05, **: p <0.01, ***: p<0.005, ****: p<0.001.

This approach was replicated in human HSCs isolated from 2 different donors and challenged with TGFβ (**Supp.** Figure 2). Of note, primary human HSCs also undergo spontaneous activation when cultured on standard cell culture plates. While TGFβ had little to no effect on *BMAL1* and *NR1D1* mRNA expression, the TGFβ-mediated overexpression of *ACTA2* and *COL1A1* was clearly decreased upon CLOCK inhibition or REV-ERB agonism at both the transcript (**Supp.** Figure 2A**, C**) and protein levels (**Supp.** Figure 2B**, D**).

Thus, these pharmacological interference experiments established the involvement of the REV-ERB/BMAL1:CLOCK axis in regulating the HSC-driven profibrotic response in both human and mouse models.

### Interaction of the REV-ERB/BMAL1:CLOCK axis with the TGFβ pathway

To further investigate the cellular mechanisms underlying the functional interplay between the TGFβ and the REV-ERB/BMAL1-CLOCK axis, we first characterized the murine HSC cell line EMS404 as a suitable model system. EMS 404 cells are SV40 large T antigen-immortalized HSCs derived from a C57Bl6 mouse (Guo et al., 2009). Upon stimulation with various profibrotic cytokines [TGFβ, angiotensin II (ATII), lipopolysaccharide (LPS), tumor necrosis factor-alpha (TNFα), and vascular endothelial growth factor (VEGF)], the transcriptional responses of CCGs and profibrotic genes to these treatments varied (**Supp.** Figure 3A). TGFβ emerged as the most consistent and potent inducer of *Serpine1*/*Pai1*, a canonical TGFβ target gene (Lund et al, 1987), along with other profibrotic genes.

HSC activation is also characterized by a metabolic switch toward aerobic glycolysis to support the energy and biosynthetic demands of activated HSCs during fibrogenesis. EMS404 cells were thus treated with TGFβ and the extracellular acidification rate (ECAR) was measured using a Seahorse extracellular flux analyzer to monitor proton (H⁺) efflux as a readout of glycolytic activity. In the presence of rotenone and antimycin A (Rot/AA), which respectively inhibit complexes I and III of the mitochondrial electron transport chain (thereby blocking mitochondrial respiration), TGFβ treatment led to a dose-dependent increase in ECAR (**Supp.** Figure 3B). The addition of 2-deoxy-D-glucose (2-DG) markedly reduced ECAR, confirming that the acidification was primarily due to glycolytic proton and lactate export. These findings indicate a robust increase in glycolytic capacity of TGFβ-activated EMS404 cells. In summary, EMS404 cells recapitulate the main metabolic and transcriptomic features of HSC transition towards an activated state.

Synchronized EMS404 cells exhibit cyclic expression of REV-ERBα, BMAL1 and CLOCK after synchronization after a serum shock (**Supp.** Figure 3C). This prompted us to develop EMS404 cells bearing a stably integrated luciferase reporter gene driven by the mouse *Bmal1* promoter (−924; +43 bp). Live recording of the bioluminescence emitted by EMS404-Bmal1Luc cells after synchronization (**Supp.** Figure 3D**, E**) yielded a cyclic pattern with a period of ≈ 22 hours in control conditions (**Supp.** Figure 3E**, F**). Interfering pharmacologically with the clock using CLK8 or SR9011 dose-dependently affected as expected this cyclic pattern (**Supp.** Figure 3E**, F**). Interestingly, neither SR9011 nor CLK8 interfered with the activation of the canonical TGFβ signaling pathway in these cells as evidenced by SMAD2&3 phosphorylation levels (**Supp.** Figure 3G) ), suggesting a direct effect of the clock further downstream of the TGFβ canonical response.

In this cellular context, TGFβ treatment shortened the Bmal1Luc period by 3.8±1.1 hours without affecting the amplitude of the bioluminescent signal (**Figure 4A**). While increasing REV-ERBα protein level with a moderate effect on cyclic *Nr1d1* mRNA levels, TGFβ increased both *Bmal1* mRNA and BMAL1 protein expression (**Figure 4B, Supp.** Figure 4A). Both CLK8 and SR9011 blunted the TGFβ-induced expression of COL1A1 at the gene and protein levels, as well as the non-rhythmic expression of *Timp1* and *Serpine1* (**Figure 4C, D**, **Supp.** Figure 4). This effect was comparable to that of SB431542, a potent blocker of the canonical TGFβ signaling pathway through TGFβ type 1 receptor (TGFBR1) kinase inhibition.

**Figure 4.**
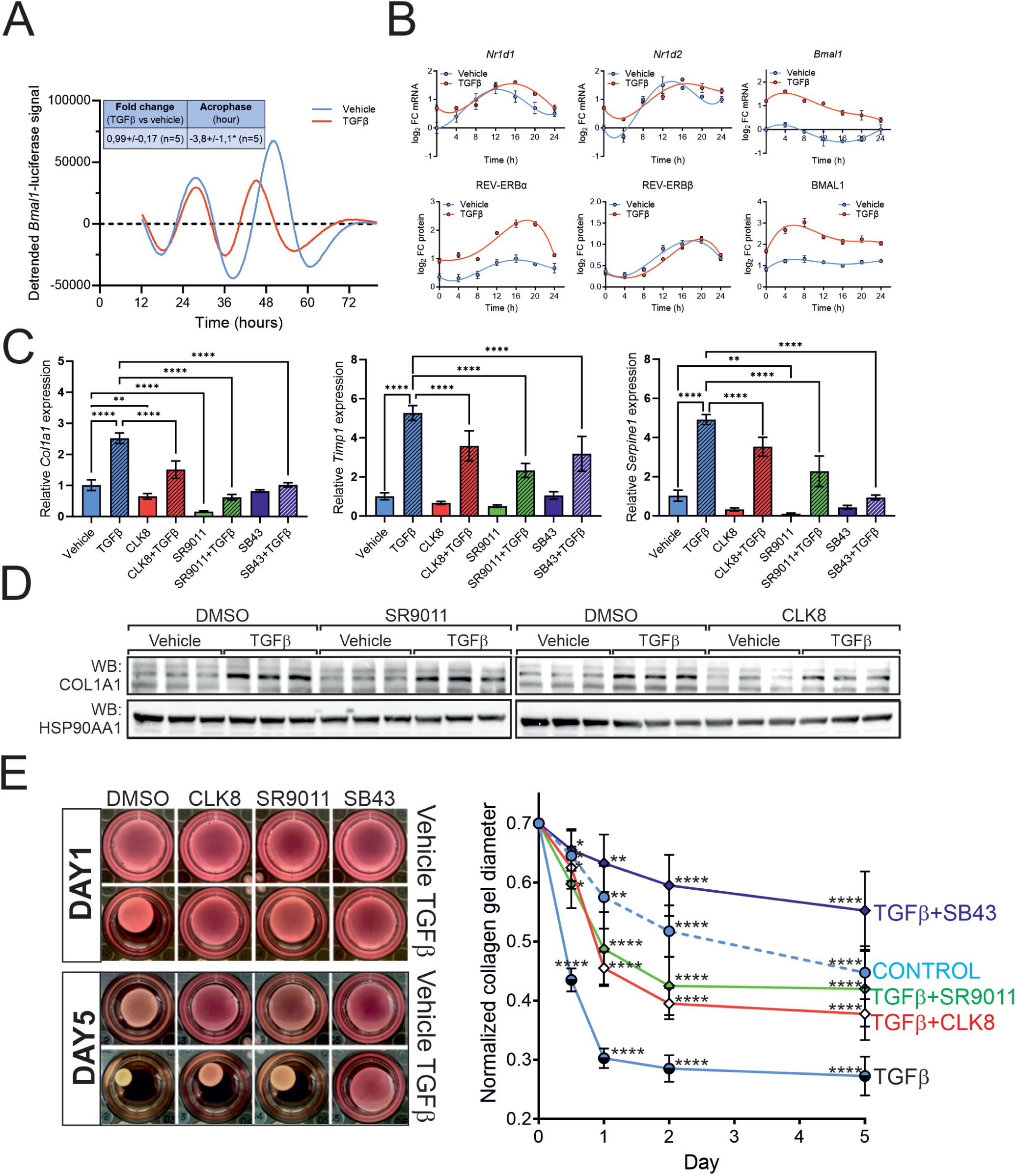
Interference with the molecular clock attenuates the fibrotic response *in vitro*. A) Bmal1-Luc rhythmicity in EMS404 cells. Synchronized EMS404 Bmal1-Luc cells were treated with vehicle or TGFβ (10 ng/mL) and the bioluminescence was recorded for over 72 h. Data were analyzed using the NiteCap software for circadian parameters (inset). B) CCG expression. Time-dependent gene (upper panel, by RT-qPCR) and protein (lower panel, by WES) expression levels were measured in synchronized EMS404 cells treated or not with TGFβ (10 ng/mL). Averaged expression values (mean ± S.E.M., n=3) are shown. C) Profibrotic gene expression in EMS cells. Profibrotic gene expression levels were measured in EMS404 cells pretreated for 24 h with indicated compounds (10 µM) followed by treatment for another 24 h with additional vehicle or TGFβ (10 ng/mL) treatment. Averaged expression values (mean ± S.E.M., n=3-5) were compared for each gene using one-way ANOVA followed by Tukey’s post hoc test. D) COL1A1 protein expression in EMS cells. EMS404 cells were treated as in C) for measurement of indicated protein levels by immunoblotting. HSP90 was used as loading control. E) Contractility assay. EMS404 cells were subjected to a collagen gel contraction assay by embedding into a solidified collagen lattice and treatment with either CLK8 (10 µM), SR9011 (10 µM) or SB431542 (1 µM) in the presence or absence of TGFβ (10 ng/mL). Representative pictures of collagen disks are shown (left panel). Daily measurements of disk diameters are shown in the right panel. Averaged expression values (mean ± S.E.M., n=3) were compared for each gene using two-way ANOVA followed by Tukey’s post hoc test. *: p<0.05, **: p <0.01, ***: p<0.005, ****: p<0.001.

Since individual gene and protein levels do not fully capture the integrated and functional cellular response to profibrotic agents, EMS404 cells were embedded in a collagen lattice to assess their contractile activity in response to TGFβ, a hallmark of hepatic stellate cell (HSC) activation. TGFβ increased the contraction of collagen disks over time, which was evaluated in the presence or absence of the test compounds SB431542, SR9011, or CLK8 (**Figure 4E**). Although less potent than the reference compound SB431242, both SR9011 and CLK8 significantly reduced TGFβ-stimulated collagen disk contraction. By the end of the linear phase (day 1), comparison of collagen disk areas showed that a TGFβ-induced 82% disk shrinkage, compared to a 32% shrinkage observed with vehicle-exposed cells, indicating both the strong pro-contractile effect of TGFβ and the baseline activation of EMS404 cells in standard 2D plastic culture. SB431542 effectively suppressed TGFβ-induced contraction, preserving 88% of the disk area, while SR9011 and CLK8 also demonstrated significant anti-contractile activity, maintaining 50% and 42% of the original disk surface, respectively.

Thus, BMAL1:CLOCK and REV-ERBα pharmacological modulation controls the fibrotic response in a representative mouse HSC cell line at both the gene expression and functional levels. These conclusions were corroborated by genetic manipulation of BMAL1 and of REV-ERBα levels either by RNA interference (BMAL1) or overexpression (REV-ERBα), which confirmed the favorable, anti-fibrotic impact of BMAL1 depletion or REV-ERB induction (**Supp Figure 5**).

**Figure 5:**
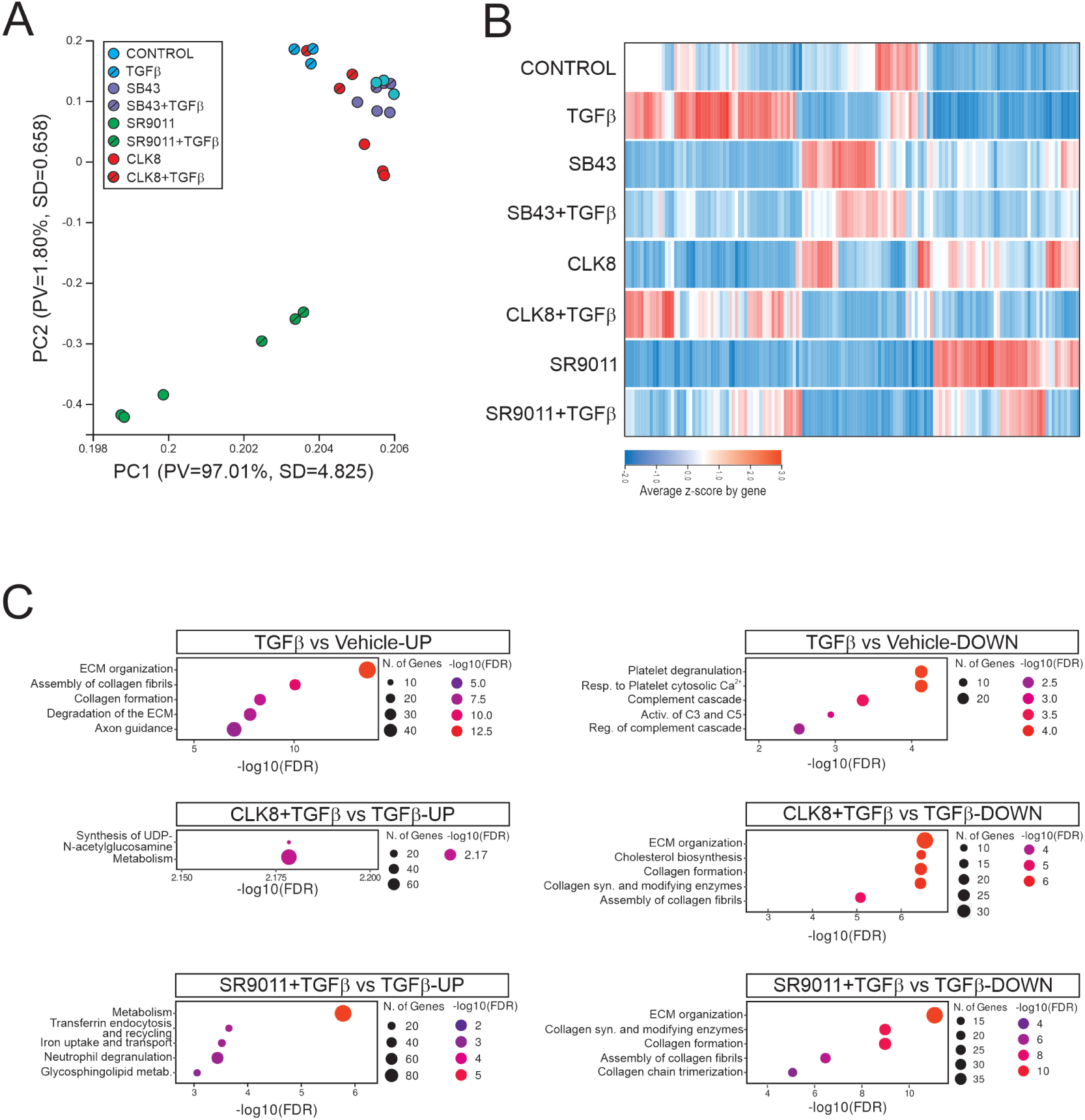
Transcriptomic analysis of the EMS404 cellular response to TGFβ and its inhibition by clock-interfering compounds. EMS404 cells were pretreated for 24 h with the indicated compound, followed by an additional 24 h of treatment with the same compound in the presence of TGFβ. A) Principal component analysis (PCA) of RNA-seq data. PCA was performed on normalized gene expression values, with each point representing an individual sample. Colors indicate the eight distinct experimental conditions. B) Heatmap of averaged gene expression values (TPM). Hierarchical clustering was applied using Euclidean distance and complete linkage. C) Biological term enrichment analysis. Following DESeq2 differential expression analysis, the top 500 up- and downregulated genes were annotated using Gene Ontology Biological Process and Reactome databases. The most significantly enriched terms are shown for each comparison.

### The REV-ERB/BMAL1:CLOCK axis essentially controls ECM-related gene expression

To identify the cellular processes triggered by TGFβ and modulated by REV-ERBs or BMAL1:CLOCK, we conducted a comprehensive RNA-seq analysis in EMS404 cells. This analysis compared untreated cells to those treated with various compound combinations (**Figure 5A**). From a quantitative perspective, TGFβ induced 1,994 transcripts (FC>1.2, q<0.05). SB431542 almost completely suppressed TGFβ-induced gene expression, repressing the expression of 1,992 (99.8%) TGFβ-induced transcripts (**Figure 5B**). Both SR9011 and CLK8 treatments exerted a less pronounced yet significant inhibitory effect on gene induction by TGFβ: 614 (31%) transcripts were fully repressed (FC<1.2) by CLK8 and 752 (37%) by SR9011 (**Supp.Table 2**). A biological term enrichment analysis revealed that TGFβ-activated ECM-related genes were significantly suppressed by both CLK8 and SR9011 (**Figure 5C**). This highlighted the potent antagonistic effects of these clock-targeting compounds on TGFβ-induced ECM remodeling, and also suggested additional, TGFβ-independent effects.

A similar analysis was applied after knocking down BMAL1 expression using siRNA as above. A biological term enrichment analysis against the KEGG database confirmed that TGFβ-upregulated genes were mostly associated to ECM-related processes (**Supp.** Figure 6A). *Bmal1* siRNA treatment markedly reduced the expression of ECM-related genes, particularly those involved in the KEGG pathways for ECM-receptor interaction and focal adhesion (**Supp.** Figure 6B-D), in line with the observed effects of CLK8 on cellular contractility (**Figure 4E**).

### Transgelin is a REV-ERBα target and regulates HSC activation in vitro

The results suggested that REV-ERB and BMAL1 modulators may share a common mechanism of action in regulating ECM-related events. We hypothesized that this effect could be driven by a shared set of gene(s) modulated by SR9011, CLK8 or the *Bmal1* siRNA. Therefore, we compared genes that were significantly upregulated or downregulated (log_2_FC > 1, q-value < 0.05) across all 3 conditions before TGFβ treatment (**Supp.** Figure 7A). Using this stringent threshold, only 28 upregulated genes and 8 downregulated genes met this criterion, thus pointing to potential key mediators underlying the convergent effects of these clock modulators on ECM-related pathways. Predicting that REV-ERBα depletion should induce an opposite effect on the expression of these key players, we performed Tandem Mass Tag (TMT)-based proteomics by mass spectrometry to identify differentially expressed proteins in naïve or REV-ERBα-depleted EMS404 cells (**Supp.** Figure 7B). This additional layer of filtering finally identified transgelin (*Tagln*) as a protein whose expression pattern aligned with responses to treatments, thus hinting at a potential role as a downstream effector.

*Tagln* is an actin-binding protein that regulates cytoskeletal dynamics and controls cell migration and contraction. Although *Tagln* is co-upregulated with other ECM-related genes such as *Col1a1*, *Acta2*, and *Dpt*, its precise functional role in HSC activation remains poorly defined. We observed that transgelin is upregulated in human fibrotic livers (fibrosis stage F > 2; **Figure 6A**) and strongly induced in HSCs during the early and later phases of CCl₄-induced liver injury in mice (**Figure 6B**). Its expression fluctuates with very low amplitude in the liver of ad libitum-fed C57Bl6/J male mice (**Figure 6C**) and was not considered as circadian, in agreement with publicly available datasets [CircaDB, (Zhang *et al*, 2014)]. *Tagln* expression is induced by TGFβ and repressed by SR9011 and CLK8 in EMS404 cells (**Figure 6D and Supp.** Figure 7A). Analysis of published ChIP-Seq data for REV-ERBα and BMAL1 in mouse liver (Annayev *et al*, 2014; Cho *et al*, 2012; Koike *et al*., 2012) detected REV-ERBα binding sites in the vicinity of the *Tagln* promoter, while no BMAL1 binding sites were detected at this locus (**Figure 6E**).

**Figure 6:**
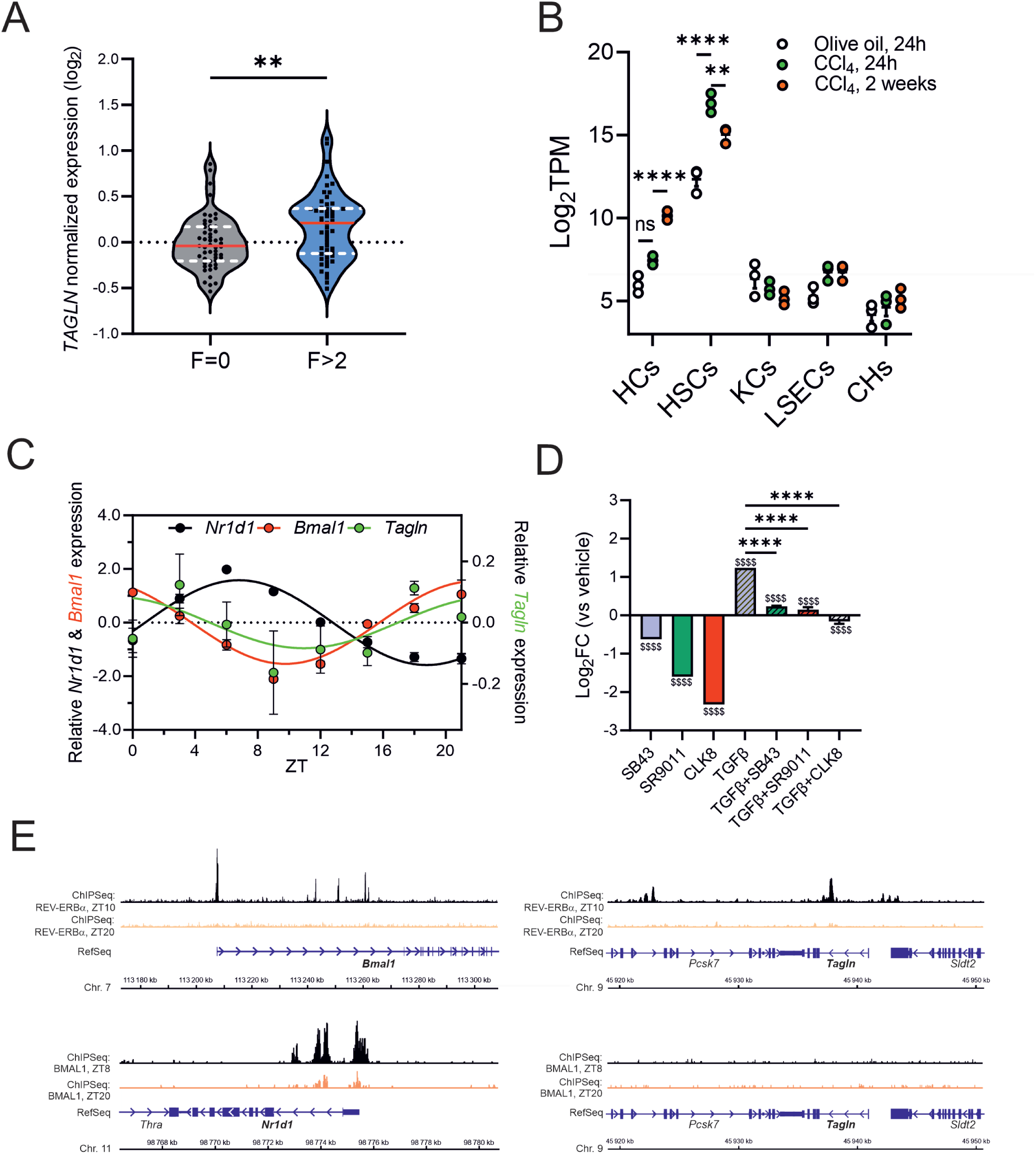
Transgelin is a clock-regulated fibrosis marker in human and mouse livers. A) Transgelin gene expression in human fibrotic livers. *TAGLN* expression values were extracted from the human fibrosis sub-cohort described in Figure 2D, encompassing 53 non-fibrotic (F=0) and 53 fibrotic (F>2) propensity-matched liver samples. Red line: median, dotted lines: 1^st^ quartile. Statistical analysis was done using an unpaired two-tailed t-test. B) Transgelin gene expression in mouse fibrotic liver. *Tagln* expression was measured by RNAseq in mouse liver cell types after CCl_4_ treatment as described in Figure 2. Averaged values (n=3, mean ± S.E.M.) were compared using two-way ANOVA followed by Tukey’s post hoc test. *: p<0.05, **: p <0.01, ***: p<0.005, ****: p<0.001. C) Time-of-day expression of *Tagln* in mouse liver. Circadian expression of *Nr1d1*, *Bmal1* and *Tagln* was determined by RT-qPCR in livers of *ad libitum*-fed mice. Data were extracted from (Zummo *et al*., 2023). D) *Tagln* expression in EMS404 cells. *Tagln* mRNA expression was measured by RT-qPCR in EMS404 cells treated with SR9011 (10 µM), CLK8 (10 µM) or SB431542 (10 µM) in the presence or absence of TGFβ (10ng/mL). Averaged values (n=3-5, mean ± S.E.M.) were compared using one-way ANOVA followed by Tukey’s post hoc test. *: p<0.05, **: p <0.01, ***: p<0.005, ****: p<0.001. E) BMAL1 and REV-ERBα genomic binding sites. REV-ERBα and BMAL1 ChIP-Seq data from bulk mouse liver were extracted from publicly available datasets (Bugge *et al*., 2012; Koike *et al*., 2012) and tracks at the *Bmal1*, *Nr1d1* and *Tagln* genomic loci were visualized using IGV.

The role of transgelin was investigated in EMS404 cells through small interfering RNA (siRNA)-mediated knockdown (**Figure 7A**). This approach efficiently abolished transgelin (*Tagln*) expression at the mRNA and protein levels, and prevented its upregulation in response to TGFβ (**Figure 7B**), known to result from phospho-SMAD2/3 binding to its cognate response element (SBE). Interestingly, *Tagln* silencing also significantly impaired TGFβ-induced *Acta2* expression under these conditions (**Figure 7C**). To evaluate whether *Tagln* and *Acta2* downregulation might affect cellular contractility, we performed a collagen gel contraction assay as above (**Figure 7D**). *Tagln* knockdown led to a ≈50% reduction in TGFβ-induced contractile activity, indicating that TAGLN significantly contributes to the generation of TGFβ-induced tension in stellate cells.

**Figure 7.**
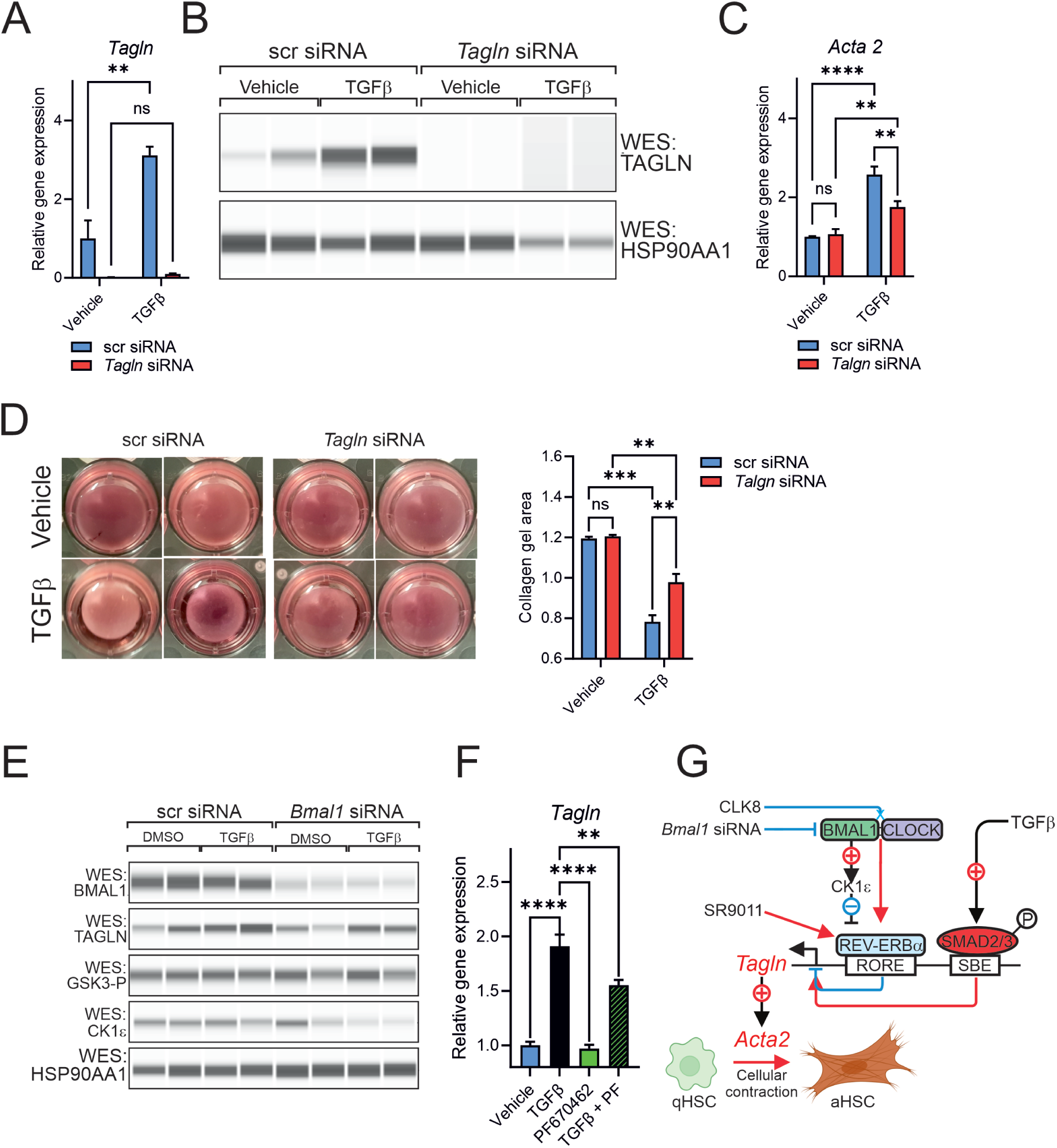
Transgelin drives HSC contractile activity. A,B) Transgelin expression in EMS404 cells. *Tagln* gene (A) and protein (B) expression was measured by RT-qPCR or WES in EMS404 cells 48 h after transfection with scrambled control or *Tagln*-targeting siRNA and treatment with TGFβ (10 ng/mL) for the last 24 h Averaged values (n=3-5, mean ± S.E.M.) were compared using two-way ANOVA followed by Tukey’s post hoc test. *: p<0.05, **: p <0.01, ***: p<0.005, ****: p<0.001. C) *Acta2* gene expression was measured as in A). Averaged values (n=3-5, mean ± S.E.M.) were compared using two-way ANOVA followed by Tukey’s post hoc test. *: p<0.05, **: p <0.01, ***: p<0.005, ****: p<0.001. D) Contraction assay. Cells were treated as in A) for a collagen gel contraction assay as in Figure 7E. Means were compared as in C). NLE) Protein expression in EMS404 cells. EMS404 cells were transfected with scrambled control or *Bmal1*-targeting siRNA and treated with vehicle or TGFβ (10 ng/mL) as in Supp. Fig. 1. Analysis of indicated proteins was performed by WES. F) Gene expression in EMS404 cells. *Tagln* expression was measured by RT-qPCR in EMS404 cells treated overnight with vehicle, TGFβ(10 ng/mL), PF670462 (10 µM) , or both compounds. Averaged values (n=3-5, mean ± S.E.M.) were compared using one-way ANOVA followed by Tukey’s post hoc test. *: p<0.05, **: p <0.01, ***: p<0.005, ****: p<0.001. G) Proposed model. A hypothetical model for the interplay between the clock and TGFβ signaling, converging on transgelin to control cell contractility, a hallmark of HSC activation, is shown. RORE: Rev-Erb/ROR Response Element; SBE: Smad Binding Element; qHSC: quiescent HSC; aHSC: activated HSC.

The comparable transcriptional effect of BMAL1(:CLOCK) inactivation and REV-ERBα agonism, combined with the lack of BMAL1 chromatin binding sites at the *Tagln* genomic locus, suggested that BMAL1 might regulate *Tagln* expression indirectly via REV-ERBα. Protein kinases are key regulators of the circadian clock by altering the stability and/or nuclear-cytoplasmic shuttling of core clock proteins (Brenna & Albrecht, 2020). Glycogen synthase kinase 3-b (GSK3β) and casein kinase 1 epsilon (CK1ε) are well-known modulators of REV-ERBα, influencing its stability and subcellular localization (Ohba & Tei, 2018; Yin *et al*, 2006). While phosphorylated GSK3β levels remained unchanged following BMAL1 depletion, CK1ε was markedly reduced under these conditions and paralleled the downregulation of TAGLN (**Figure 7E**). Based on the hypothesis that CK1ε may limit REV-ERBα nuclear localization, hence its transrepressive activity, we treated EMS404 cells with the dual CK1δ/ε inhibitor PF670462 (Choi *et al*, 2023) and assessed *Tagln* expression (**Figure 7F**). PF670462 significantly decreased expression in presence of TGFβ, mimicking the effects of BMAL1 inhibition.

Taken together, these observations indicate that TAGLN is a key regulator of stellate cell contractility via a BMAL1-CK1ε-REV-ERBα axis, the precise molecular mechanisms of which remain to be elucidated.

## DISCUSSION

The molecular clock plays a central role in liver physiology and pathology. Its function as a key regulator of lipid and glucose metabolism, as well as redox pathways, has been extensively studied. The consequences of clock disruption on these processes have been characterized in both mouse models and human studies (Daniels *et al*, 2023). Given this regulatory capacity, the causative relationship between circadian disruption and the progression of MASLD toward MASH and hepatic fibrosis has fostered intense scrutiny. Notably, it was suggested that components of the molecular clock may represent promising pharmacological targets for chronic liver diseases (Crouchet *et al*., 2025; Ni *et al*, 2023). However, the extent to which the circadian clock is disrupted in human liver disease, whether this disruption affects non-parenchymal liver cell types and contributes to disease progression remains unclear. Previous findings from our group demonstrated that core clock genes (CCGs) retain differential expression between morning and afternoon liver biopsies in human MASH and fibrosis (Johanns *et al*., 2024). Using multiparametric propensity score matching, accounting for sex, BMI, diabetes status, and statin use, we show here for the first time that CCG mRNAs daily oscillations are altered in human MASH and fibrotic livers (**Figure 1, Supp.** Figure 1). Interestingly, these alterations are also detected in steatotic livers, indicating that metabolic dysregulations impinge on the molecular clock. This aligns well with reported dysregulations of circadian rhythmicity in livers from high fat-fed mice (Abbondante *et al*, 2016; Eckel-Mahan *et al*, 2013; Kohsaka *et al*, 2007). While informative about human pathology, the use of liver biopsies does not allow for distinguishing between different liver cell types, as the transcriptomic signal is likely dominated by hepatocytes.

This question was investigated using the pericentral hepatotoxicant CCl_4_ in mice from which major liver cell types were separated by cell sorting. This showed that CCG expression is sensitive to chemical liver injury, mostly in HCs and HSCs. Conversely, altering circadian rhythmicity through pharmacological manipulation of CLOCK:BMAL1 or REV-ERBα coincided with decreased expression of TGFβ-induced HSC activation markers in mouse liver explants and human primary HSCs (**Figures 3, 4, Supp.** Figure 2). This is in line with studies in pulmonary and liver fibrosis showing beneficial effects of REV-ERBα agonism (Crouchet *et al*., 2025; Cunningham *et al*, 2020; Prasad *et al*, 2023; Wang *et al*, 2023).

Whole body knockout of *Bmal1* reduced liver fibrosis markers in surviving, leptin-deficient mice; however, it is unclear whether this effect is due to beneficial pleiotropic metabolic changes or results from cell-autonomous mechanisms (Jouffe *et al*., 2022). Along this line, BMAL1 overexpression in TGFβ-stimulated human hepatic stellate LX2 cells inhibited glycolysis, proliferation, and phenotypic activation (Xu *et al*., 2022). In our study, REV-ERBα activation and BMAL1:CLOCK inactivation limited the transition from a quiescent to an activated phenotype, these processes were molecularly characterized in the well-defined mouse hepatic stellate cell line EMS404. Both REV-ERBα agonism and BMAL1:CLOCK blockade significantly blunted TGFβ-induced profibrotic gene expression and cellular contractility, a hallmark of HSC activation (**Figure 4**). Transcriptional profiling of SR9011 or CLK8-treated EMS404 cells revealed their clear TGFβ-antagonizing effect, characterized by a downregulation of regulons associated with ECM remodeling and cellular migration. This pattern mirrored the effects observed with *Bmal1* siRNA treatment, ruling out nonspecific effects of CLK8. Interestingly, primary HSCs from mice expressing the dominant-negative mutant of CLOCK ClockΔ19 showed accelerated activation in vitro (Jokl *et al*., 2023), a finding consistent with our observations. However, the CLOCK-controlled regulon showed only very minor overlap with our REV-ERBα- and Bmal1 siRNA/CLK8 regulon. The data rather pointed to Rho GDP-dissociation inhibitors as potential molecular relays of CLOCKΔ19-mediated defects. In fact, we also noted that Rho GTPase signaling components contribute to profibrotic gene expression and alter circadian rhythmicity in EMS404 cells (data not shown). This highlights the reciprocal relationship between ECM architecture, the cytoskeleton and the clock machinery which remains to be fully investigated (Dudek *et al*, 2023). It also suggests that core clock components probably act through distinct signaling pathways to affect the same biological processes.

The mechanistic links between circadian dysregulation and HSC activation remain poorly defined. Through integrated transcriptomic and proteomic analyses following treatment with various clock perturbagens, transgelin was identified as a key relay of the BMAL1-REV-ERBα and TGFβ signaling axis in HSCs. *Tagln* expression is SMAD-dependent in TGFβ-activated HSCs (Gnainsky *et al*, 2007; Yu *et al*, 2008) and, according to single cell data, is induced in CCl_4_- and bile duct ligation-activated HSCs (Liu *et al*, 2020; Yang *et al*, 2021). Its expression correlates with ECM-related genes in multiple cell types (Weiskirchen, 2023) and its inhibition has been proposed as a valuable strategy to maintain physiological functions in lung and kidney fibrosis (Kwon *et al*, 2024; Yu *et al*., 2008). *Tagln* knockdown blunted *Acta2* induction by TGFβ in our model (**Figure 7**). Although limited, this observation suggests that the roles of transgelin extend beyond being a mere effector of cellular contractility via its interaction with actin filaments. Indeed, emerging evidence suggests it also plays regulatory roles by engaging with signaling pathways such as the PARP–Rho axis, potentially relaying cytoskeletal cues to the nucleus and influencing gene expression (Rasmussen & Jin, 2024).

Taken together, our findings provide strong evidence for a dysregulated yet functionally active circadian clock in both human and mouse models of hepatic fibrosis. Pharmacological targeting of core clock components, such as REV-ERBα and BMAL1, effectively limited HSC activation. This effect appears to involve the downstream effector TAGLN, at which clock and TGFβ signaling pathways converge, as illustrated in our synthetic model (**Figure 7G**). REV-ERBα-mediated repression of *Tagln* expression is dependent on BMAL1-driven induction of CK1ε, a kinase known to phosphorylate phosphodegrons in several clock proteins including REV-ERBα (Narasimamurthy & Virshup, 2021). Here, we provide new insight into how clock-controlled signaling integrates with broader regulatory networks, such as TGFβ, to govern HSC activation. Whether this mechanism extends to additional molecular mediators of HSC activation remains to be explored. It also paves the way for exploring novel treatment options for liver fibrosis, and potentially beyond.

## ACKNOWLEDGEMENTS

This work was supported by ANR (LABX EGID, ANR-10-LABX-0046). MJ was supported by grants from Wallonie-Bruxelles International (WBI, Belgium, ref. SUB/2020/479801) and European Association for the Study of the Liver (EASL, Sheila Sherlock fellowship). GT is supported by an ANR grant (ANR-23-CE14-0028-01). This work was supported by a grant from ANR to PL (ClockFib, ANR-23-CE14-0028-01). PL’s and BS’s teams were supported by grants from Fondation pour la Recherche Médicale (EQU202203014645 and EQU202203014650, respectively).

## Author contributions

Conceptualization: MJ,AB,PL; Methodology: MJ,AB,JV,GT,DV,GH; Software: JV,JDC; Validation: MJ,AB,JV, MV,FPZ,DV,BS; Formal analysis: MJ,AB,JV,JDC,DV,GH,PL; Investigation: MJ,AB,SC,MV,NV,FPZ,MBG,GTCG,GH,VR,FP; Resources: MJ,AB,FPZ,MBG,VR,FP,BS,JE; Data Curation: AB,MJ,JDC,JV,PL; Writing: MJ,AB,BS,JE,PL; Visualization: MJ,AB,JV,JDC,SC,MV,FPZ,GT,DV,PL; Supervision: PL; Project administration: MJ,PL; Funding acquisition: MJ,BS,PL.

---

**Supplemental Figure 1:**
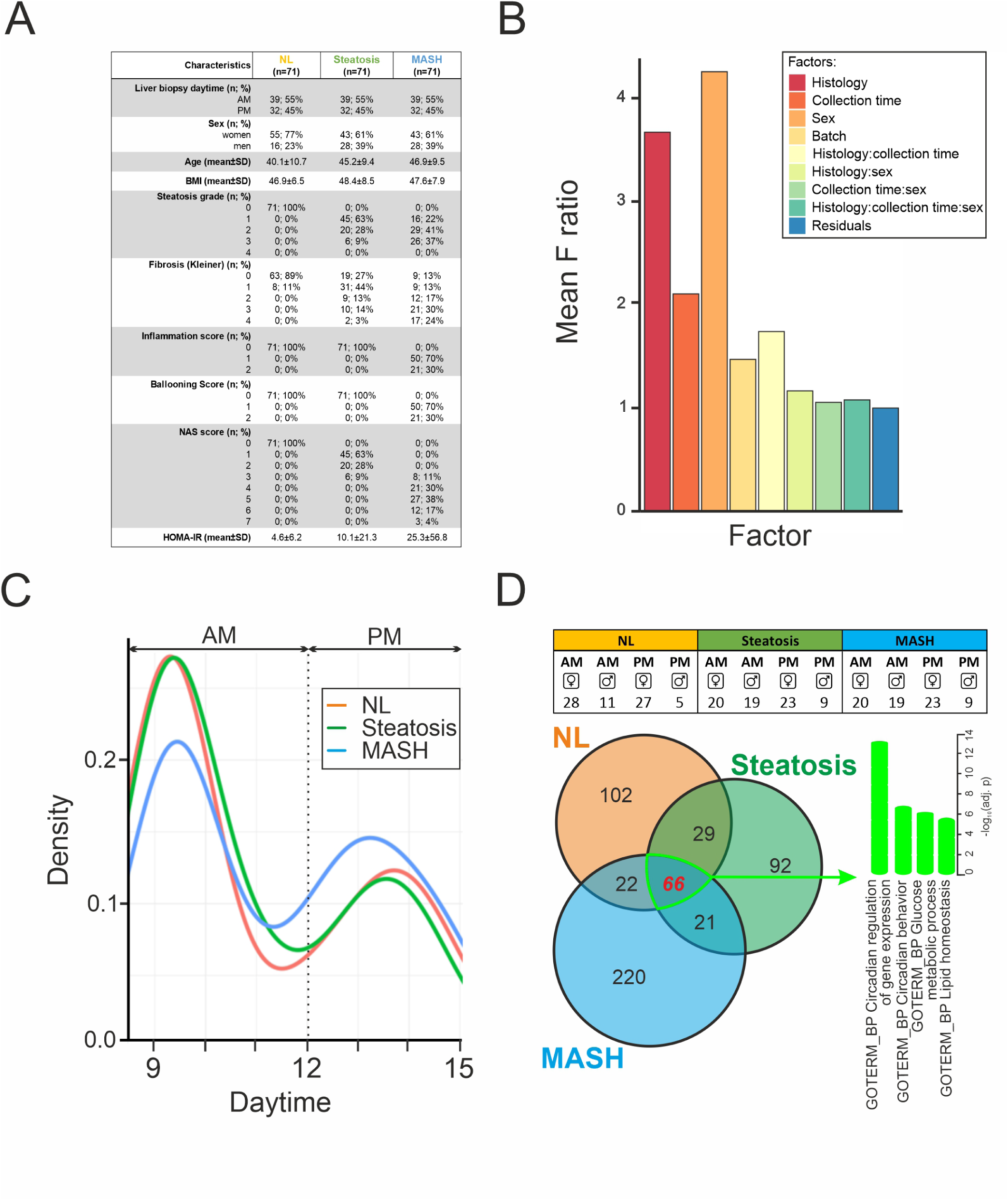
Daytime-dependent gene expression in a human MASH cohort. A) The propensity score-matched cohort. Biometric, biochemical, and histological parameters of propensity score-matched patients from whom liver biopsies were taken at various daytimes and histologically classified as normal (NL; n=71), steatotic without pathology (n=71) or affected by metabolism-associated steatohepatitis (MASH; n=71). Continuous values were expressed as mean ± SD. Inter-group comparisons were performed using the Wilcoxon test for continuous variables and the exact Fisher test for categorical variables. AM: ante meridiem, PM: post meridiem, BMI: body-mass index, NAS: NAFLD activity score, HOMA-IR: homeostatic model assessment of insulin resistance. B) Sources of variation in liver biopsies. ANOVA was used to identify sources of variation between gene expression profiles of liver biopsy samples. C) Relative frequency distribution of liver biopsy collection daytimes between NL, steatosis and MASH groups. D) Overlap of daytime-dependent genes identified by RNA-seq in normal, steatotic or MASH livers. The upper panel indicates the group distribution of the sex of patients, a known confounding factor.

**Supplemental Figure 2.**
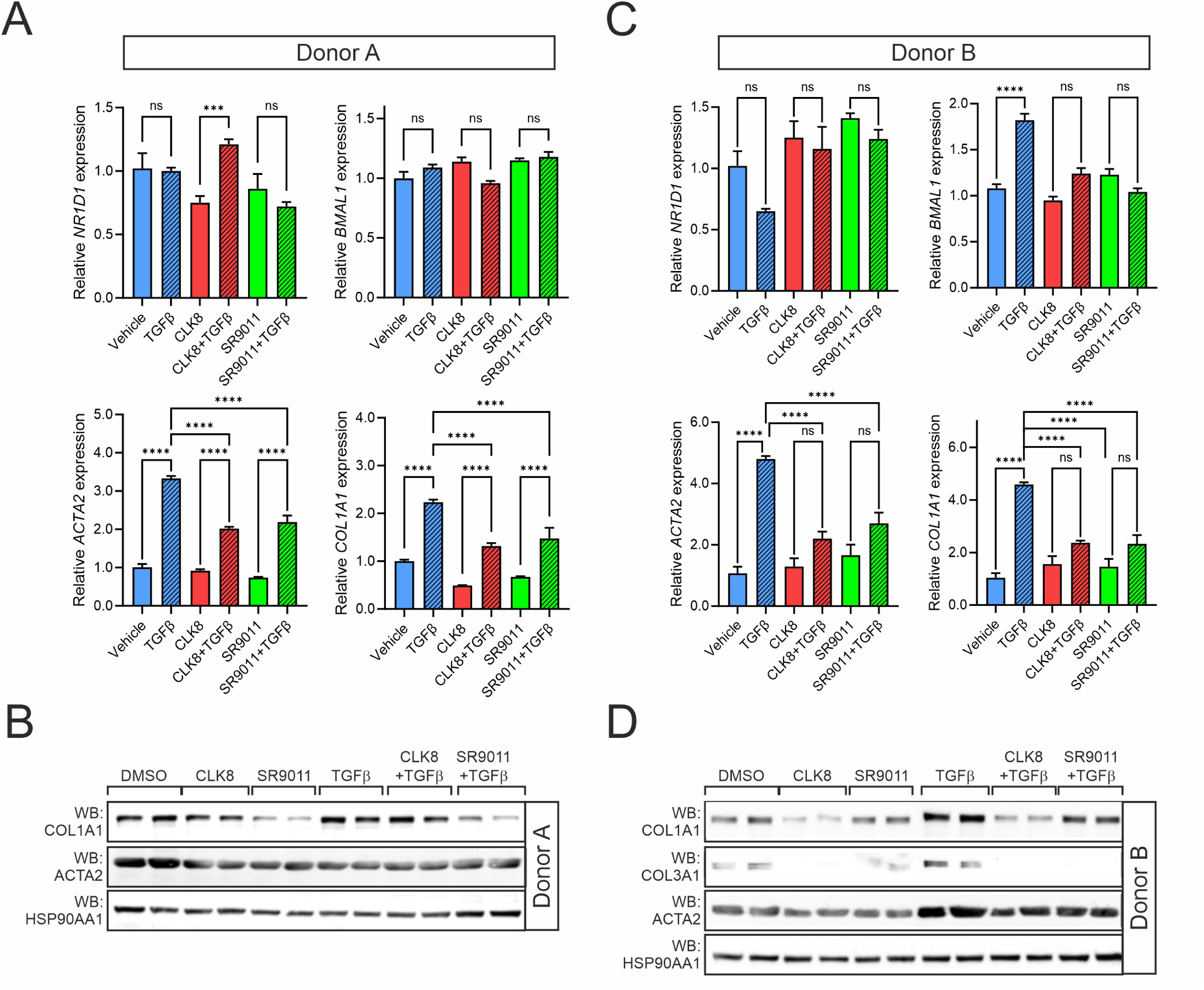
C**l**ock **perturbagens and the response to TGF**β **of human primary HSCs.** Primary HSCs were isolated from 2 different donors (A, B: donor A; C, D: donor B). A,C) Gene expression levels. Expression levels of selected clock genes (upper panels) and fibrosis markers (lower panels) were measured by RT-qPCR in cells pretreated for 24 h with the indicated compounds (5 µM), followed by an additional 24 h in the presence of TGFβ (10 ng/mL) added on top. Averaged expression values (mean ± S.E.M., n=3) were compared for each gene using one-way ANOVA followed by Tukey’s post hoc test. *: p<0.05, **: p <0.01, ***: p<0.005, ****: p<0.001. B, D) Profibrotic protein expression. Cell extracts were obtained after incubation in the same conditions as in A) and analyzed by immunoblotting using the indicated antibodies against fibrosis markers or HSP90 as a loading control.

**Supplemental Figure 3.**
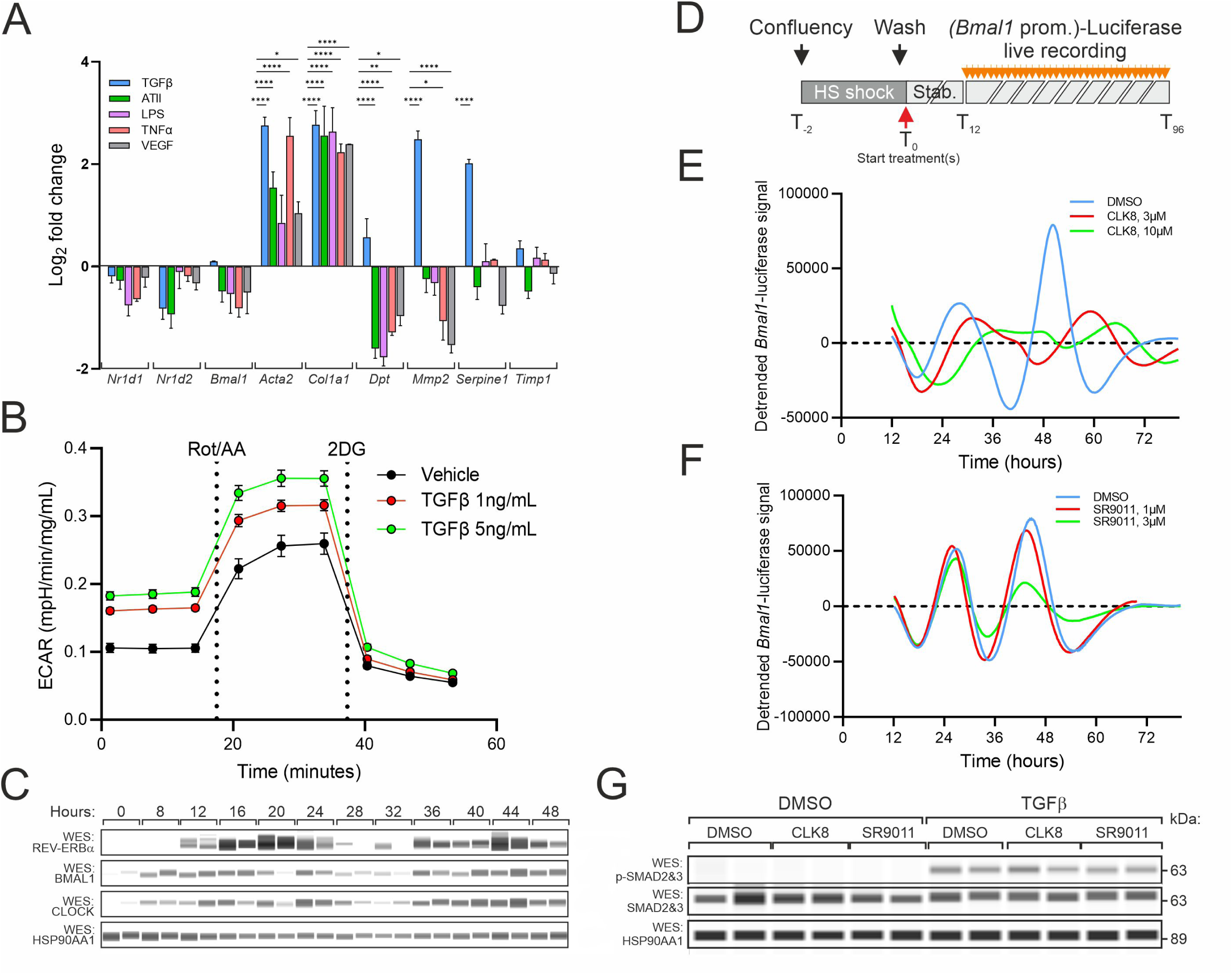
T**h**e **mouse cell line EMS404 as a model for HSC activation and clock cycling.** A) EMS404 cellular sensitivity to chemo/cytokines. EMS 404 cells were treated for 24 h with different profibrotic stimuli: TGFβ (10 ng/mL), angiotensin II (ATII, 10 µM), lipopolysaccharide (LPS, 100 ng/mL), tumor necrosis factor α (TNFα, 10 ng/mL), or vascular endothelial growth factor (VEGF, 20 ng/mL). Expression of clock and fibrosis marker genes was analyzed by RT-qPCR. Results were expressed as fold changes relative to untreated conditions. Averaged expression values (mean ± S.E.M., n=3) were compared for each gene using one-way ANOVA followed by Tukey’s post hoc test. *: p<0.05, **: p <0.01, ***: p<0.005, ****: p<0.001. B) Glycolytic activity of EMS404 cells. Extracellular acidification rate (ECAR) of EMS404 cells in response to TGFβ was measured using a Seahorse XF analyzer at the basal state in presence of glucose and after addition of rotenone/antimycin A (Rot/AA) followed by 2-deoxyglucose (2-DG). C) CCG protein levels. REV-ERBα, BMAL1 and CLOCK protein expression in synchronized EMS404 cells was analyzed by the WES capillary-based immunoassay using the indicated antibodies. D) Experimental outline for circadian rhythm analysis in Bmal1-Luc–expressing EMS404 cells. Cells were synchronized by a serum shock (2 h with horse serum, HS) followed by a stabilization (Stab.) period of 12 h before continuous monitoring of bioluminescence for 4 days. E, F) Rhythmicity activity of the Bmal1-Luc transgene in the presence of the CLOCK:BMAL1 disruptor CLK8 or of the REV-ERBα agonist SR9011. G) The canonical TGF pathway in EMS404 cells. Activation of the canonical TGFβ-pSMAD2/3 pathway was assessed by WES in extracts of EMS404 cells treated with TGFβ (10 ng/mL) for 20 min after pretreatment for 24 h with CLK8 (5 µM) or SR9011 (5 µM).

**Supplemental Figure 4:**
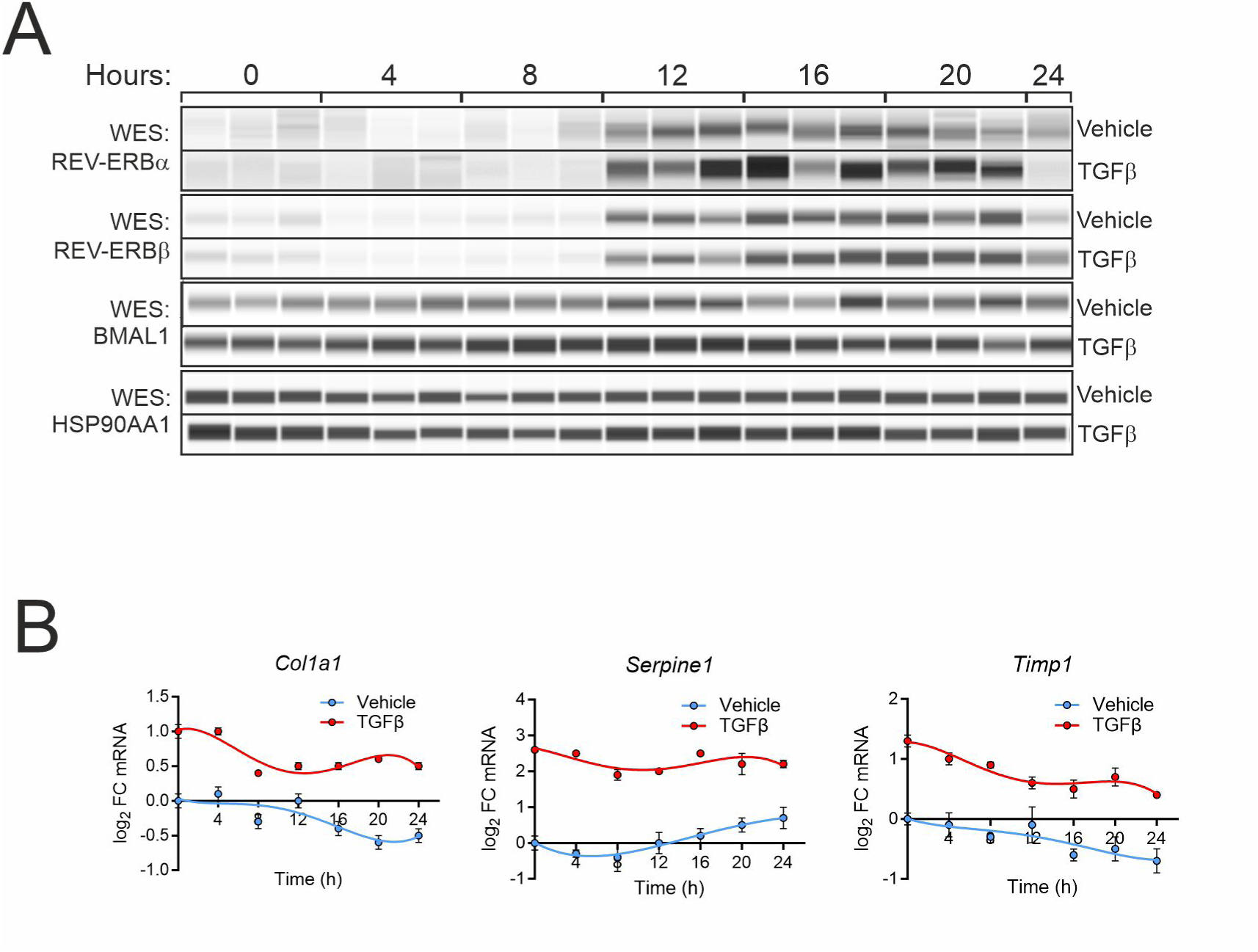
**Effect of TGF**β **on circadian expression of clock components and fibrosis markers.** EMS404 cells were synchronized and treated with vehicle or TGFβ (10 ng/mL). Cells were collected at 4 h intervals for protein extraction and analysis by WES with indicated antibodies (A) or mRNA extraction and quantification by RT-qPCR (B). The quantitative analysis of these data is shown in Figure 4B.

**Supplemental Figure 5.**
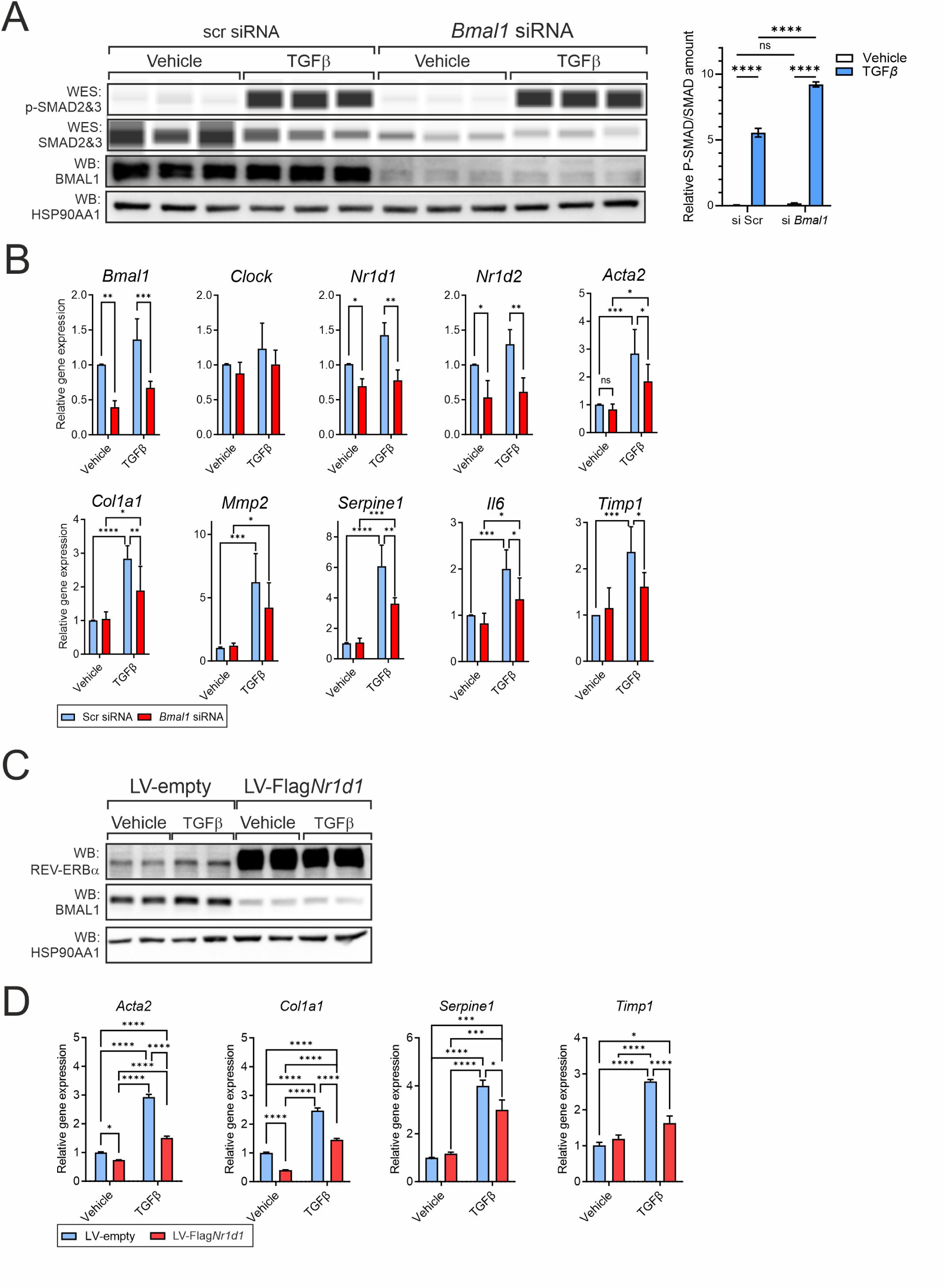
G**e**netically **targeting BMAL1 affects TGFβ signaling and downstream effects.** A) The canonical TGF pathway in BMAL1-depleted EMS404 cells. EMS404 cells were transfected with scrambled control, siRNA, or a pool of siRNAs targeting *Bmal1* transcripts. After 48 h, cells were exposed to vehicle or TGFβ (10 ng/mL) for 20 min. Protein levels were assessed by capillary-based WES immunoassay, and P-SMAD/total SMAD quantification is shown in the panel on the right. B) Gene expression in BMAL1-depleted EMS404 cells. EMS404 cells were treated as in A), except that TGFβ was added for the last 24 h, and mRNA content was assayed by RT-qPCR. Averaged expression values (mean ± S.E.M., n=3) were compared for each gene using an two-sided ANOVA. *: p<0.05, **: p <0.01, ***: p<0.005, ****: p<0.001. C, D) REV-ERBα overexpression in EMS404 cells. Flag-tagged REV-ERBα was stably overexpressed in EMS404 cells using a lentivirus (LV-Flag*Nr1d1*) compared to an empty control vector (LV-empty). Puromycin-selected cell pools were treated with vehicle or TGFβ (10 ng/mL) for 24 h for protein extraction and analysis by WES (C) or mRNA extraction and quantification by RT-qPCR (D). Averaged expression values (mean ± S.E.M., n=3) were compared for each gene using an two-sided ANOVA. *: p<0.05, **: p <0.01, ***: p<0.005, ****: p<0.001.

**Supplemental Figure 6:**
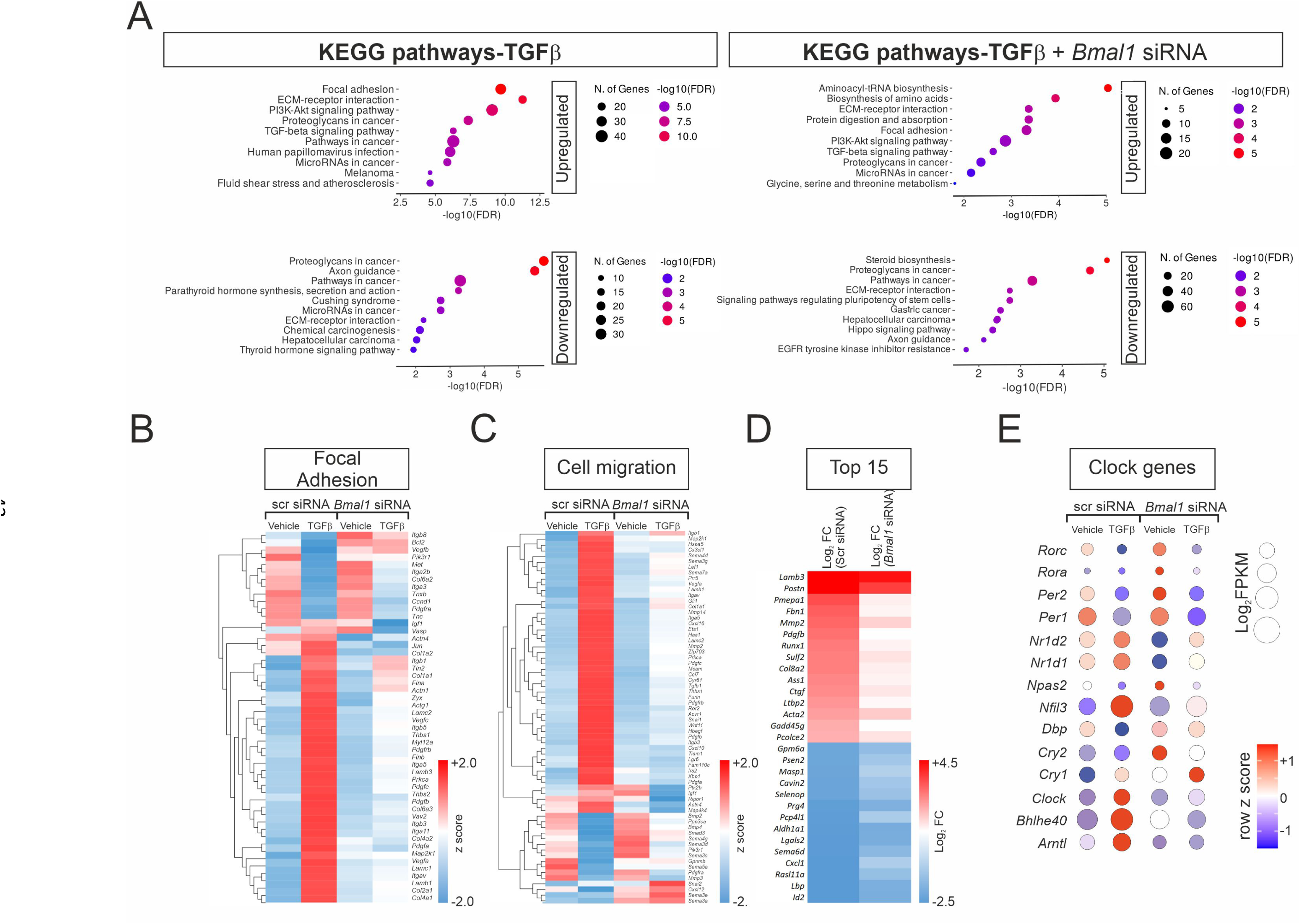
Effect of BMAL1 silencing on the transcriptomic response to TGFβ. EMS404 cells were transfected with a scrambled control or *Bmal1*-targeting siRNA pool and treated 24 h later with vehicle or TGFβ (10 ng/mL). Gene expression was measured by RNAseq and analyzed using DESeq2. A) BTEA in in BMAL1-depleted EMS404 cells. A biological term enrichment analysis was carried out using the KEGG database on the top 500 TGFβ-induced genes in the presence or the absence of BMAL1. The most significantly enriched terms are shown for each comparison. Heatmaps show group-averaged gene expression values (TPM) for genes related to the terms focal adhesion (B), cell migration (C) and the top 15 up- and downregulated genes (D) in BMAL1-depleted EMS404 cells. E) RNAseq expression data of CCGs in control and BMAL1-depleted EMS404 cells treated with or without TGFβ.

**Supplemental Figure 7:**
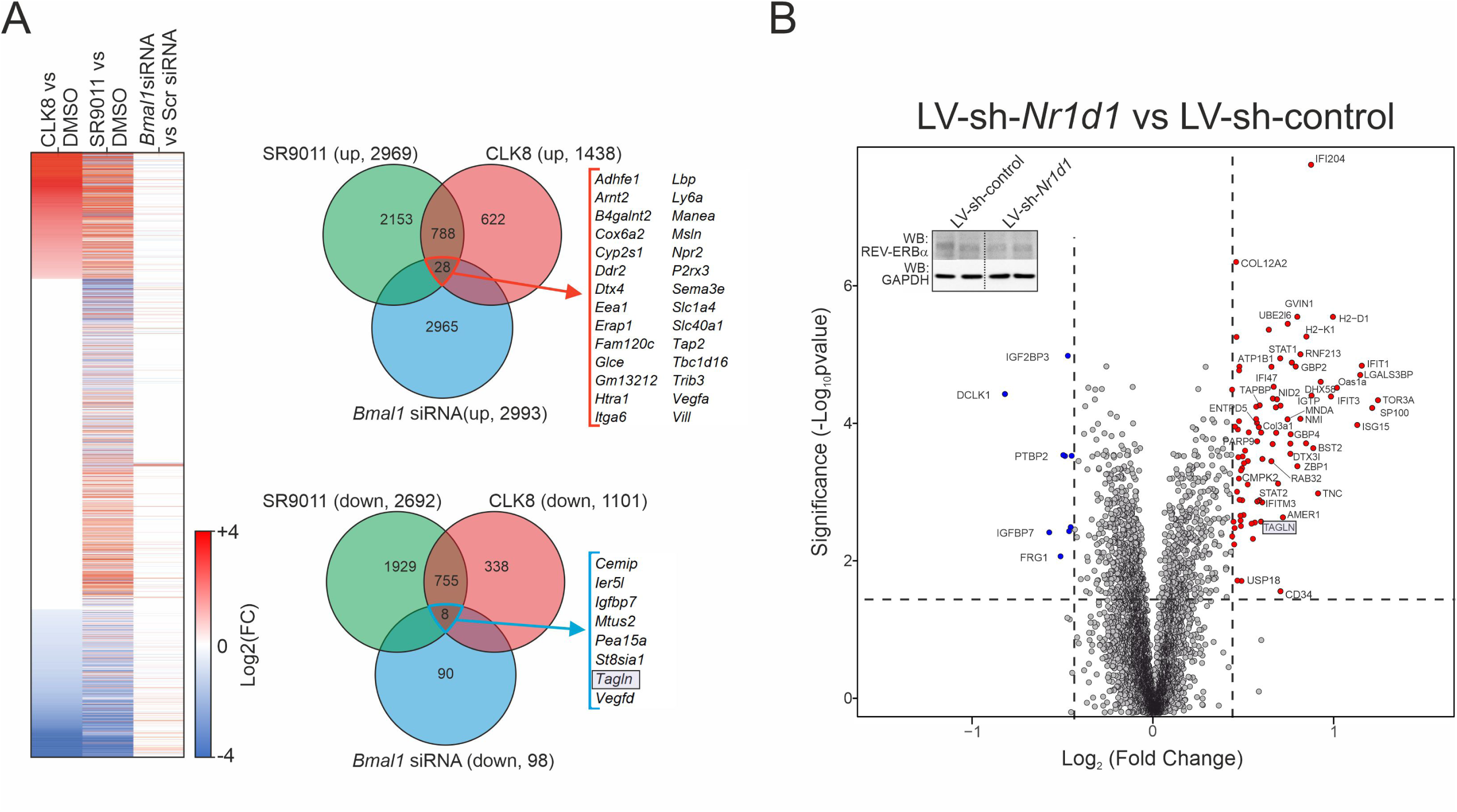
Identification of mediators for clock-regulated modulators of fibrosis. A) Alteration of the EMS404 cell basal gene expression pattern by clock perturbagens. EMS404 cell transcriptomic data were analyzed for overlapping up- or downregulated genes (|FC|>1.2, q<0.05 between indicated conditions). Left panel: Heatmap of group-averaged gene expression fold changes. Hierarchical clustering was applied using Euclidean distance and complete linkage. Right panel: Venn diagram indicates the number of differentially expressed genes between conditions. B) Mass spectrometry analysis of proteins in REV-ERBα-depleted EMS404 cells. EMS404 cells were lentivirally (LV) transduced with a short hairpin RNA targeting REV-ERBα (LV-sh*Nr1d1*) or a control shRNA. Intracellular proteins were analyzed by comparative mass spectrometry using a TMT labelling-based quantification approach. Transgelin (*Tagln*/TAGLN) is highlighted as a common hit.

